# A choroid plexus apocrine secretion mechanism shapes CSF proteome and embryonic brain development

**DOI:** 10.1101/2024.01.08.574486

**Authors:** Ya’el Courtney, Joshua P. Head, Elizabeth D. Yimer, Neil Dani, Frederick B. Shipley, Towia A. Libermann, Maria K. Lehtinen

## Abstract

We discovered that apocrine secretion by embryonic choroid plexus (ChP) epithelial cells contributes to the cerebrospinal fluid (CSF) proteome and influences brain development in mice. The apocrine response relies on sustained intracellular calcium signaling and calpain-mediated cytoskeletal remodeling. It rapidly alters the embryonic CSF proteome, activating neural progenitors lining the brain’s ventricles. Supraphysiological apocrine secretion induced during mouse development by maternal administration of a serotonergic 5HT2C receptor agonist dysregulates offspring cerebral cortical development, alters the fate of CSF-contacting neural progenitors, and ultimately changes adult social behaviors. Critically, exposure to maternal illness or to the psychedelic drug LSD during pregnancy also overactivates the ChP, inducing excessive secretion. Collectively, our findings demonstrate a new mechanism by which maternal exposure to diverse stressors disrupts *in utero* brain development.

## INTRODUCTION

The epithelial cells of the choroid plexus (ChP), located in each brain ventricle, are a principal source of cerebrospinal fluid (CSF) (*1*, *2*) as well as secreted factors that play important roles in the central nervous system throughout life (*3–5*). For example, disrupting temporal regulation of CSF-borne signals during embryonic development leads to impaired neural progenitor proliferation and brain abnormalities that persist into adulthood (*6*, *7*). ChP epithelial cells introduce elements such as proteins, lipids, mRNA, and miRNA into the CSF via vesicles and exosomes (*8–11*). Nonetheless, the basic mechanisms by which ChP epithelial cells regulate CSF signal availability are poorly understood. Gaining insight into embryonic ChP secretion will improve our comprehension of brain development and highlight potential developmental threats.

Calcium signaling represents a broadly conserved mechanism by which cells regulate secretion (*12*, *13*). ChP epithelial cells exhibit spontaneous, transient calcium (Ca^2+^) activity, with the potential for larger, prolonged increases in response to stimuli like serotonin (5-HT) (*14*). Among serotonergic receptors, 5HT2C (*Htr2c*, a G_q_/G_11_-coupled G protein-coupled receptor) is the most highly expressed by ChP epithelial cells (*15*). Activating ChP 5HT2C is accompanied by Transferrin (*16*) and Insulin (*17*) release, though the mechanisms of release have not been characterized. We recently demonstrated in adult mice that ChP 5HT2C can be activated by selective synthetic agonist WAY-161503 (*18*), leading to a rise in intracellular Ca^2+^, expression of activity-dependent genes, and *in vivo* observations of apocrine secretion in adult mice (*19*). Similar to the ChP, secretory epithelial cells in salivary glands, mammary glands, coagulating glands, and sweat glands are equipped for vesicular and exosomal secretion, but notably also utilize apocrine secretion (*20–23*), a high-capacity release mechanism that introduces intracellular contents into the extracellular space. Collectively, these observations motivated the present study to characterize ChP apocrine secretion and interrogate its physiological importance in shaping the developing CSF proteome.

## RESULTS

### Embryonic ChP epithelial cells introduce proteins into the CSF via apocrine secretion

Embryonic ChP is equipped for and uses an activity-dependent apocrine mechanism to secrete factors into the CSF that coordinate brain development. To test our hypothesis that ChP apocrine secretion mediates release of CSF proteins that signal to neural progenitor cells and impact cortical development, we first assessed whether apocrine machinery functions at timepoints relevant to cerebral cortical neurogenesis. In mice, neurogenesis primarily occurs from embryonic day 10.5 (E10.5) to E17.5 (*24*) (**Fig. 1A**). The lateral ventricle (LV) ChP, nearest the ventricular zone where these progenitors reside, begins to differentiate around E9.5 (*3*). In mice, 5HT2C is highly and selectively expressed in ChP epithelial cells from E12 to E18.5 (**Fig. 1B, Fig. S1**) (*25–27*). Remarkably, maternal subcutaneous administration of the 5HT2C agonist WAY-161503 caused potent activation of ChP at E12.5, as evaluated by activity-dependent c-Fos protein expression (**Fig. 1C-D**). We identified ChP apocrine structures by scanning electron microscopy (SEM), transmission electron microscopy, H&E staining, and immunohistochemistry (**Fig. 1E**). They were fairly homogenous, arising singly from a depression in the center of the apical membrane, immediately adjacent to the ciliary bundle (**Fig. S2A-C, Movie S1**) supported by a network of microtubules (**Fig. S2D, Movie S2**) and protruded an average of 5µm from the cell surface (**Fig. S2E**). Maternal administration of WAY-161503 continued to activate the ChP at E14.5 and E16.5, evaluated by activity-dependent gene transcription (**Fig. 1F**) and apocrine secretion in approximately 70% of embryonic ChP epithelial cells (**Fig. 1G**, quantified in **Fig. 1H**). We used SEM to visualize the appearance and distribution of apocrine structures in the LV ChP of adult mice (**Fig. 1I**) and found a match between the embryonic and adult apocrine secretion phenomena.

**Figure 1.**
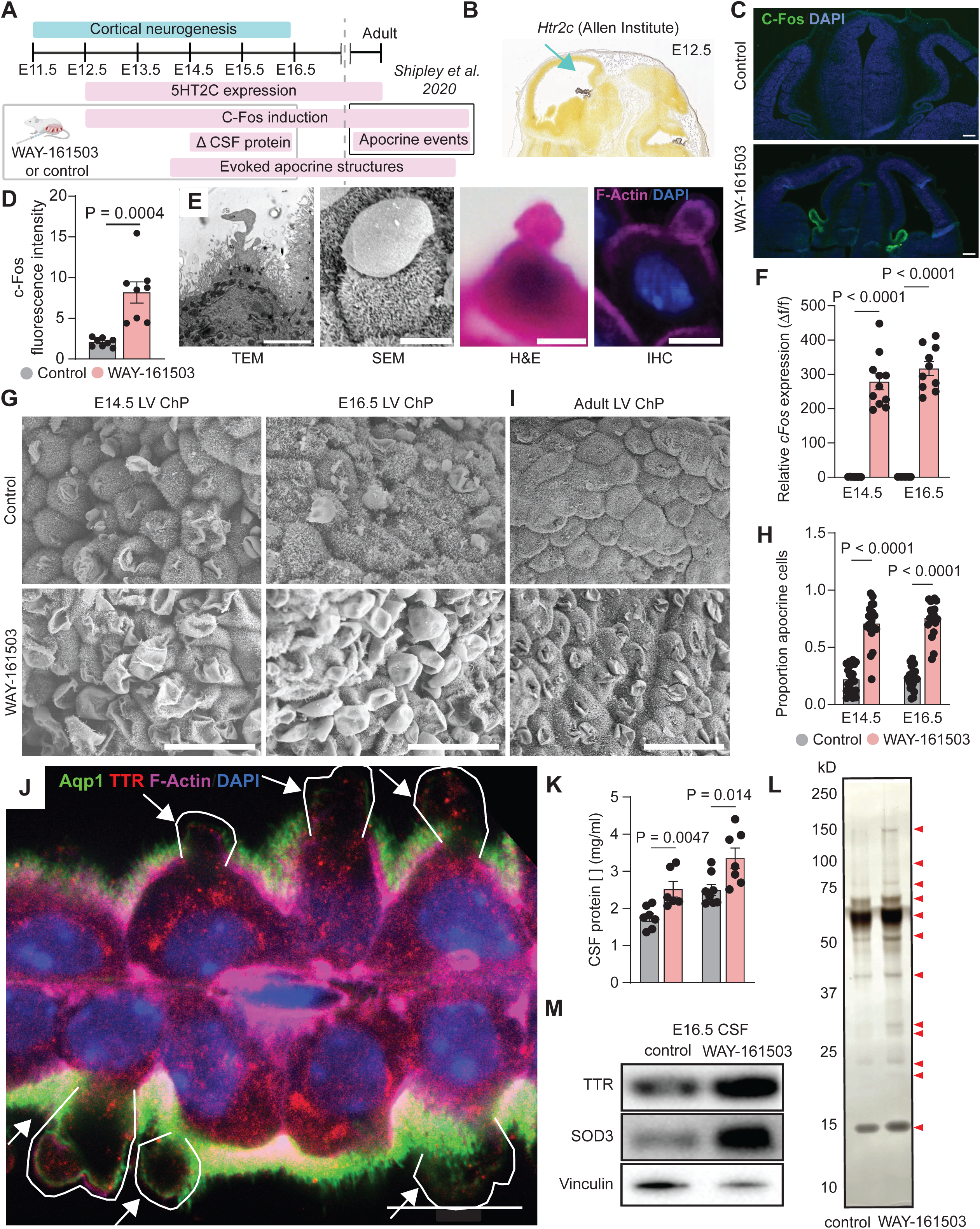
Activating embryonic ChP 5HT2C evokes apocrine secretion of proteins into CSF. (**A**) Timeline of readouts of apocrine secretion in relation to cortical development timepoints. (**B**) *Htr2c* expression in E12.5 mouse brain. Allen Developing Mouse Brain Atlas, developingmouse.brain-map.org/experiment/show/100047619. (**C**) Representative confocal images of E12.5 mouse brain 30’ after maternal saline (control) or WAY-161503 injection. Scale bars 100µm. (**D**) Intensity of c-Fos immunofluorescence in E12.5 ChP, *n* = 8 embryos from 4 litters/condition. P values from Mann-Whitney test. (**E**) Apocrine structures visualized by, from left to right, TEM, SEM, H&E stain, and IHC. Scale bars 5µm. (**F**) RT-PCR for ChP *cFos* at E14.5 (left, *n* = 11 embryos, 3 litters) and E16.5 (right, *n* = 10 embryos, 3 litters). P values from one-way ANOVA. **(G)** SEM of E14.5 (left) and E16.5 (right) LV ChP. Scale bars = 20µm. (**H**) Quantification of (G). *N* = 20 embryos, 4 litters/condition, P values from Mann-Whitney test. (**I**) SEM of adult LV ChP. Scale bars 40µm. (**J**) ProExM of E16.5 LV ChP 30’ after maternal WAY-161503 injection. Scale bar 20µm. (**K**) Embryonic CSF protein concentration 30’ after maternal saline (control) or WAY-161503. Each point represents pooled CSF from one litter. E14.5 control *n* = 7 WAY *n* = 6, E15.5 control *n* = 4 WAY *n* = 6, E16.5 control *n* = 8 WAY *n* = 7. P values from Mann-Whitney test. (**L**) Silver stain of E16.5 CSF 30’ after maternal saline (control, left) or WAY-161503 (right). Red arrows indicate proteins whose concentration differed between conditions. (**M**) Western blot for TTR and SOD3. All data presented as means ± SEM.

Protein-retention expansion microscopy (*28*) and immunohistochemistry revealed that apocrine structures contained important cargo including transthyretin (TTR), a signature protein of the ChP (**Fig. 1J**). BCA assays (**Fig. 1K**) and silver staining (**Fig. 1L**) of CSF collected at baseline or following apocrine stimulation revealed that apocrine secretion was associated with increased presence of many proteins. Targeted approaches confirmed increased availability of TTR and the antioxidant superoxide dismutase 3 (SOD3) (**Fig. 1M**). Our data cannot rule out the possibility that these proteins could also be released by other modes of secretion, as 5HT2C signaling is also accompanied by ChP vesicle fusion events (*19*). In addition to proteins and as described in other epithelia (*22*), organelles including mitochondria were observed in apocrine structures (**Fig. S2E**).

Programmed cell death (apoptosis) is a common event that results in the apical extrusion of unhealthy cells from epithelial sheets (*29*). To ensure that the structures we identified as apocrine were not actually apoptotic bodies, we performed cleaved Caspase-3 staining and evaluated nuclear chromatin condensation but did not observe any evidence of apoptosis after evoked apocrine secretion (**Fig. S3**). Furthermore, the apoptotic bodies described in other epithelia are irregularly shaped, and often found in cells with multiple protrusions and compromised structural integrity and polarity (*29–31*). By contrast, the structures we identified as apocrine arose as a single protrusion per cell, and each cell’s structural integrity and polarity were maintained below the apical membrane (**Fig. 1J**, **Fig. S2D**).

### Embryonic ChP apocrine secretion releases CSF signaling factors and activates proliferative pathways

To determine potential biological functions of embryonic apocrine secretion, we conducted an untargeted analysis of proteins in E16.5 CSF at baseline vs. 30 minutes after WAY-induced apocrine secretion using the SomaScan v4.1 aptamer-based proteomics platform. Of 7,596 proteins represented on the platform, 6,561 were detected in mouse CSF, and 123 proteins were differentially detected in CSF following maternal WAY-161503 exposure (false discovery rate-adjusted P<0.05, **Table S1, Data S1**).

Of the 123 CSF proteins that were differentially abundant following apocrine secretion, 101 increased in abundance and 22 decreased (**Fig. 2A**). Among those increased were two notable proteins with neurodevelopmental relevance: Insulin-like growth factor 2 (Igf2) and Sonic hedgehog (Shh) (**Fig. S4A**). CSF-Igf2 stimulates progenitor cell proliferation and is required for healthy cortical development (*6*) whereas CSF-Shh is important for proper brain size and hindbrain development (*32*). Using a ChP explant-conditioned medium approach (**Fig. 2B**), we validated release of both Igf2 and Shh following WAY-161503 treatment. Importantly, the ChP tissues are patterned, with ventricle-specific transcriptomes and secretomes (*33*). *Htr2c* (**Fig. S4B**) and *Igf2* (**fig. S4C**) expression were consistent across ventricles, and we observed similar Igf2 release into conditioned medium by lateral and 4^th^ ventricle ChP (**Fig. 2C**). For proteins known to have regional expression (**Fig. S4D-E**), we confirmed Shh release selectively by 4^th^ ventricle ChP (**Fig. 2D**) (*33*) and highest insulin release by 3^rd^ ventricle ChP (**Fig. 2E**) (*17*).

**Figure 2.**
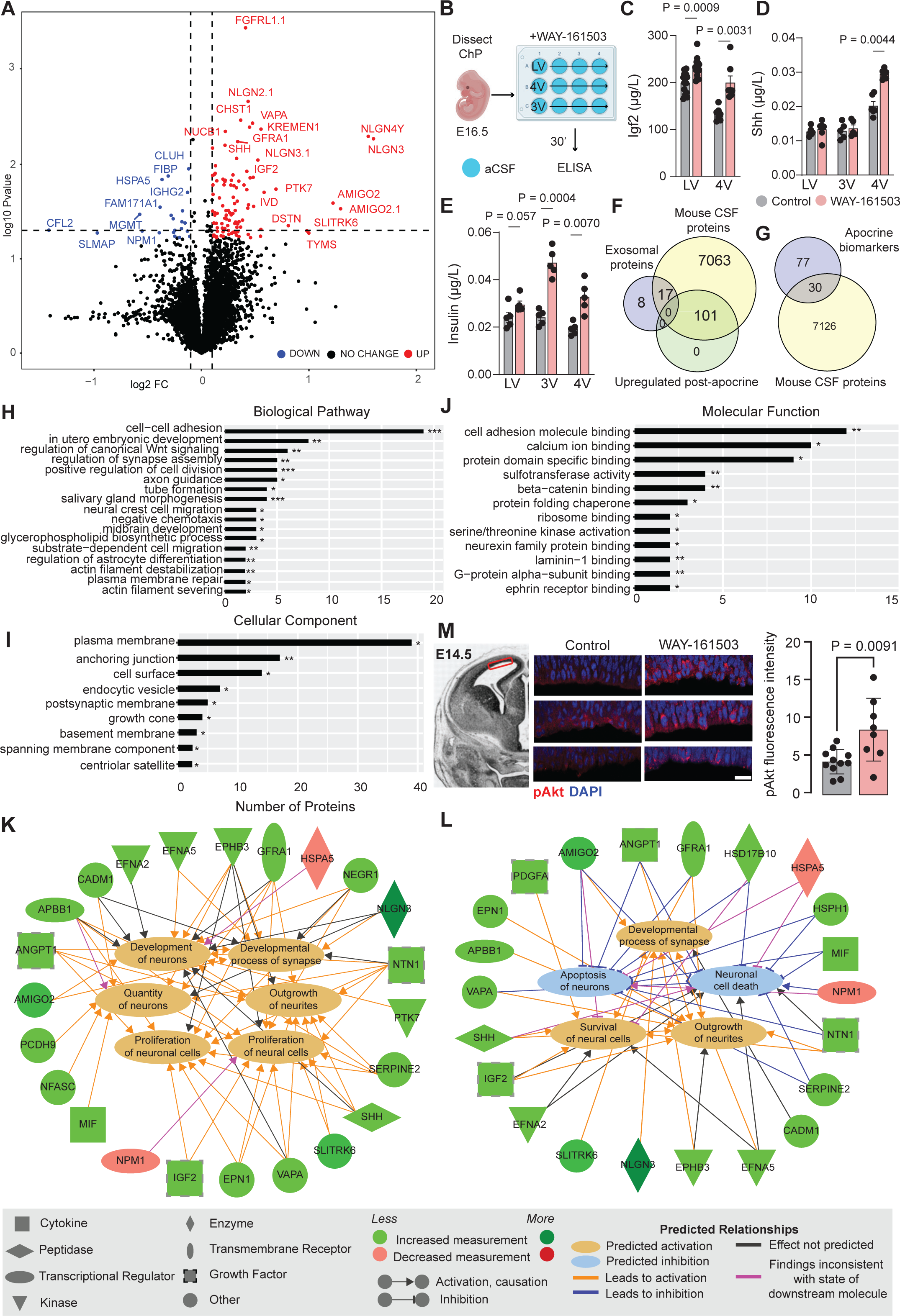
Apocrine secretion alters the embryonic CSF proteome. (**A**) Schematic of embryonic CSF collection for proteomics. (**B**) Volcano plot showing differently abundant proteins between E16.5 CSF at baseline and following apocrine secretion. Proteins with *p-*values less than 0.05 appear above the horizontal dashed line. The vertical dashed lines indicate the threshold of ±1.1-fold change. *N* = 3 pooled litters/condition. (**C**) Experimental schematic for data presented in (D) and (E). (**D**) ELISA of explant-conditioned aCSF for IGF-2. Each point represents aCSF conditioned by 5 E16.5 ChP explants. LV *n* = 14, 3V *n* = 7, 4V *n* = 7. P values from Mann-Whitney test. (**E**) ELISA of explant-conditioned aCSF for SHH. Each point represents aCSF conditioned by 5 E16.5 ChP explants. LV *n* = 6, 3V *n* = 5, 4V *n* = 6. P values from Mann-Whitney test. (**F**) Venn diagram shows exosomal proteins present in baseline CSF but not released by apocrine secretion. (**G**) Venn diagram shows overlap between CSF proteins and biomarkers of apocrine secretion. (**H**) Significant biological pathways identified using gene set enrichment analysis (GSEA). For (H) as well as (I) and (J), statistical significance was determined as an FDR-corrected P < 0.05 (one-sided) using right-tailed Fisher’s exact test. (**I**) Significant cellular components identified using GSEA. (**J**) Significant molecular functions identified using GSEA. (**K**) Ingenuity pathway analysis of apocrine-secreted proteins demonstrated activation of proliferation- and development-related pathways and (**L**) inhibition of cell-death pathways. (**M**) Confocal images of pAkt in E14.5 neural progenitor cells (left) and quantification of pAkt fluorescence intensity (right). Control *n* = 11, 3 litters, WAY-161503 *n* = 8, 3 litters. P values from Mann-Whitney test. All data presented as means ± SEM.

To determine whether 5HT2C activation also induces release of extracellular vesicles (EV), we tested the intersection between baseline CSF proteins, post-apocrine CSF proteins, and a list of 25 often-identified EV markers (*34*) (**Fig. 2F**, EV marker list in **Table S2**). As expected, 17 of 25 EV markers were present in control CSF. However, no EV markers increased in CSF following 5HT2C activation (**fig. S4F**), suggesting that the contents of apocrine secretion do not include membrane-bound EVs. To firmly place ChP epithelial cells among apocrine tissues, we assessed the intersection between baseline CSF proteins and a list of 122 proteins considered as biomarkers of apocrine secreted fluids (*35*) (**Fig. 2G**). Embryonic CSF contained 77 of the 122 proteins (**Table S3**) that are conserved in all apocrine fluids, regardless of anatomical origin, lending further support to our model that apocrine secretion occurs in ChP epithelial cells and can be evoked by 5HT2C activation.

To obtain a comprehensive understanding of the breadth of CSF contents arising from apocrine secretion, we used Gene Ontology (GO) enrichment analysis and the Ingenuity Pathway Analysis (IPA) software tool (QIAGEN, Redwood City, CA) (*36*) to reveal interactions between differentially abundant CSF proteins and their potential downstream targets. GO enrichment analysis revealed an over-representation of several biological pathway terms with implications for neural development, including regulation of Wnt signaling, cell proliferation, differentiation, and migration, as well as neuronal functions including regulation of synapse assembly, axon guidance, and neural crest cell migration (**Fig. 2H**). Overrepresented cellular component terms were consistent with an apocrine mechanism and included plasma membrane, anchoring junction, and cell surface proteins (**Fig. 2I**). Overrepresented molecular functions were also associated with apocrine secretion (*22*), including cell-adhesion molecule binding and calcium ion binding (**Fig. 2J**). IPA revealed networks of apocrine protein interactions supporting neuron quantity and proliferation (**Fig. 2K**), activating neural survival pathways, and inhibiting neuronal cell death and apoptosis pathways (**Fig. 2L**). Taken together, these analyses support the model that embryonic ChP apocrine secretion not only provides a CSF niche favorable to cell viability, but also signals likely to facilitate healthy proliferation, differentiation, and migration of neural progenitor cells in the ventricular zone. Indeed, we found that E14.5 ventricular zone neural progenitor cells had increased phosphorylated AKT 30 minutes following WAY-161503 evoked secretion (**Fig. 2M**), suggesting that altered CSF contents stimulated downstream signaling events in target cells.

### Embryonic overstimulation of apocrine secretion disrupts cortical development

Since apocrine secretion altered the embryonic CSF proteome and stimulated downstream signaling events in neural progenitors, we sought to understand whether evoking supraphysiological levels of apocrine secretion in embryos would affect cerebral cortical development. We chronically overstimulated apocrine secretion by administering WAY-161503 subcutaneously to pregnant dams daily from E12.5 to E16.5. After the pups were born, we evaluated the distribution of excitatory neurons marked by Ctip2, Satb2, and Tbr1 at postnatal day 8 (P8) and gross brain morphology at P30 (**Fig. 3A**). While we failed to see evidence of gross differences in brain morphology or overall cell density at P30 (**Fig. S5**), we observed increased numbers of Satb2+ neurons and fewer Tbr1+ neurons in the primary somatosensory cortex (**Fig. 3B**). Across all cortical layers, numbers of Satb2+ neurons increased by 7.9% ± 2.6%, Tbr1+ neurons decreased by 10.3% ± 2.6%, and Ctip2+ neurons were not affected. We then divided the cortex into six equal-sized bins to ascertain the relationship between these cell population changes and migration patterns. This analysis revealed that population changes were driven by alterations in the deeper cortical layers reflected in bins 4, 5, and 6. Satb2+ populations increased by 14.2% in bin 4, 13.8% in bin 5, and 15.9% in bin 6 (**Fig. 3C**). Tbr1+ populations decreased by 19.1% in bin 5 and 27% in bin 6 (**Fig. 3D**). These data suggest that overstimulating apocrine secretion in otherwise healthy pregnant mice alters cortical excitatory neuronal populations, shifting the identity of a proportion of deep-layer neurons from Tbr1+ corticothalamic projection neurons to Satb2+ callosal projection neurons. This underscores the sensitivity of apical progenitors in the ventricular zone—the CSF-contacting progenitor cells that divide earlier in neurogenesis and form the deeper cortical layers (*37*, *38*),—to CSF composition.

**Figure 3.**
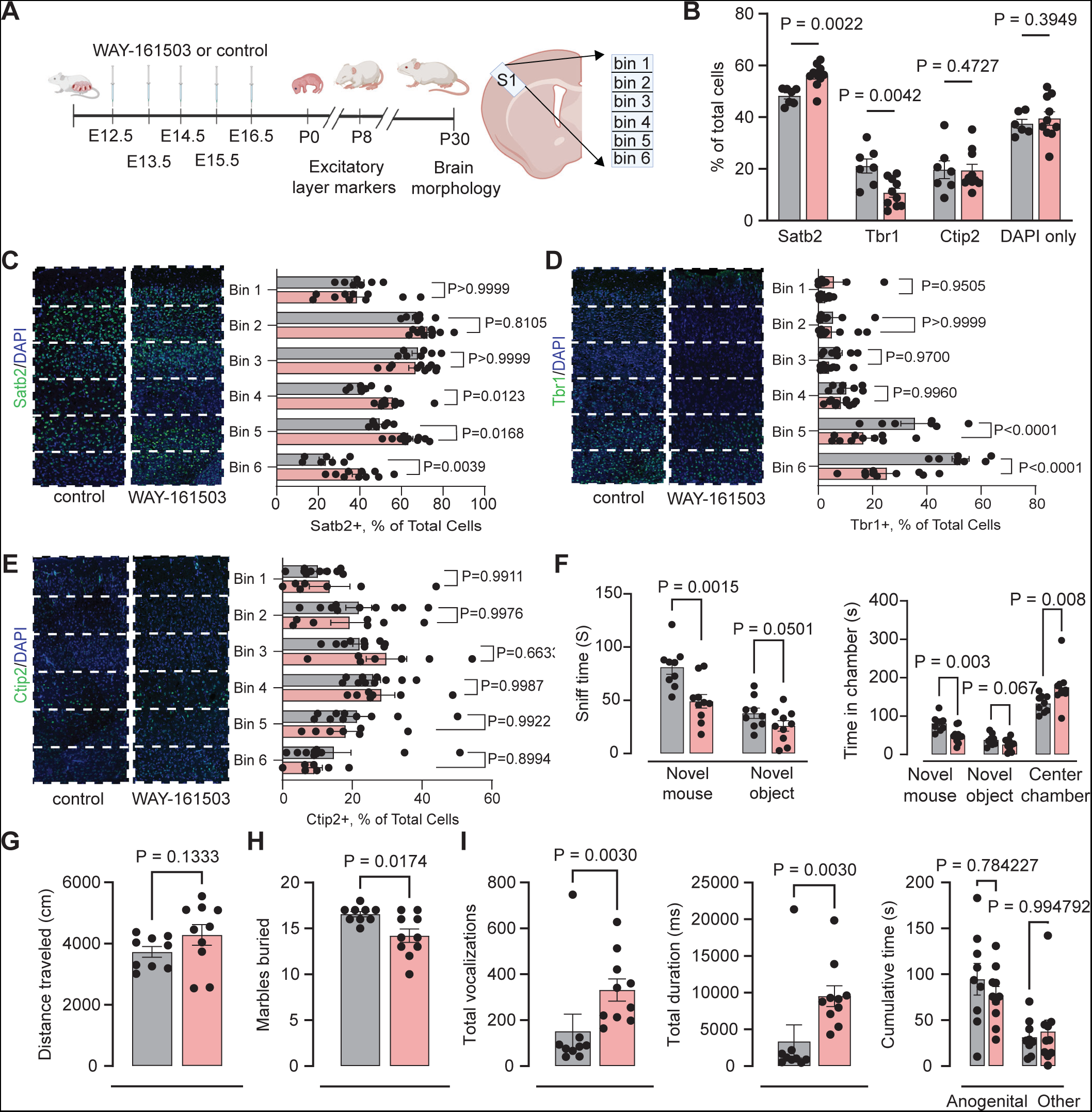
Chronic embryonic overstimulation of apocrine secretion leads to abnormal cerebral cortical development and adult behavior. (**A**) Schematic depicting experimental timeline and readouts. (**B**) Quantification of Ctip2+, Tbr1+, and Satb2+ neuronal populations in P8 S1 cortex in control (*n* = 18 mice, 4 litters) and chronic apocrine (*n* = 20 mice, 4 litters) mice. P values from nonparametric Kruskal-Wallis test. (**C**) (left) Confocal images of Satb2 (green) in S1 cortex. (right) Satb2+ neuron proportion by bin, P values from Mann-Whitney test. (**D**) (left) Confocal images of Tbr1 (green) in S1 cortex. (right) Tbr1+ neuron proportion by bin, P values from Mann-Whitney test. (**E**) (left) Confocal images of Ctip2 (green) in S1 cortex. (right) Ctip2+ neuron proportion by bin, P values from Mann-Whitney test. (**F**) Three-chambered social approach task using *n* = 9 control, *n* = 10 apocrine mice. (left) Time spent directly sniffing novel mouse or object. (right) Time in chamber with novel mouse, novel object, or in center. (**G**) Distance travelled (in cm). P values from Mann-Whitney test. (**H**) Graph of marbles buried (*n* = 9 control, *n* = 10 apocrine). P values from Mann-Whitney test. (**I**) Male-female ultrasonic vocalizations (*n* = 9 control, *n =* 10 apocrine). (left) Total vocalizations. (center) Total duration (ms) of vocalizations. (right) Cumulative time (s) that a male mouse spent in physical contact with the female mouse. P values from Mann-Whitney test. All data presented as means ± SEM.

These changes in cerebral cortical cytoarchitecture were accompanied by several behavioral phenotypes in adult offspring. Mice that experienced supraphysiological embryonic apocrine secretion were allowed to mature to P90, then three literature-validated (*39*, *40*) behavioral assays for assessing neurodevelopmental disorder-associated phenotypes were administered: three-chamber social approach (*41*), marble burying (*42*), and adult male-female ultrasonic vocalizations (USVs) (*43*). Abnormal behavior arose in the three-chamber social approach task, where chronic apocrine mice displayed lower sociability than control mice and spent less time in the chamber with or interacting with the novel mouse (**Fig. 3F**). Distance traveled (**Fig. 3G**) and velocity (**Fig. S6A**) were unchanged, suggesting that decreased time spent with the novel mouse was not attributable to motor deficits. Chronic apocrine mice also buried fewer marbles in the marble-burying task (**Fig. 3L**), an established metric for assessing rodent repetitive behaviors. Marble burying was impacted by sex as a biological variable, with male mice burying fewer marbles and no change in female mice (**Fig. S6B**). Differences attributable to sex were not significant in any other analyses performed, either cortically or behaviorally. In the USV task, chronic apocrine mice were significantly more vocal, emitting more total courtship vocalizations that lasted for longer, but did not initiate more physical interactions with the test mouse (**Fig. 3I**). When USVs were broken down into more nuanced subcategories based on frequency and pattern, this effect appeared to be a nonspecific increase across most types of vocalizations (**Fig. S6C**).

### Diverse maternal stimuli evoke embryonic apocrine secretion via a convergent mechanism

Curious to know if more diverse stimuli affect neurodevelopment through the ChP-CSF system, we tested and found several additional physiological scenarios including psychedelics, temperature-sensitive receptor activation, and maternal illness trigger apocrine secretion. Psychedelic compounds, including lysergic acid diethylamide (LSD), act strongly at 5HT2C (*44*) and have been reported to act in adult ChP (*45*). Maternal exposure to a translationally relevant dose of LSD (*46*, *47*) (0.2 mg/kg) resulted in transmission of the drug to the developing embryonic brain, as evidenced by ChP activation at E16.5. 30 minutes following subcutaneous administration of LSD to pregnant dams, we observed that LSD reached the embryonic brain, activating ChP immediate early gene transcription (**Fig. 4A**) and apocrine secretion (**Fig. 4B**). C-Fos expression increased by 70-fold compared to control, which while smaller than the 380-fold increase observed with WAY-161503 (**Fig. 4C, Fig. S7A**), was still associated with sufficient activation to cause apocrine secretion. We observed apocrine structures in 68% of E16.5 ChP epithelial cells after LSD exposure, a 46% increase from baseline and similar to the 78% of cells exhibiting apocrine structures after WAY-161503 (**Fig. 4D**).

**Figure 4.**
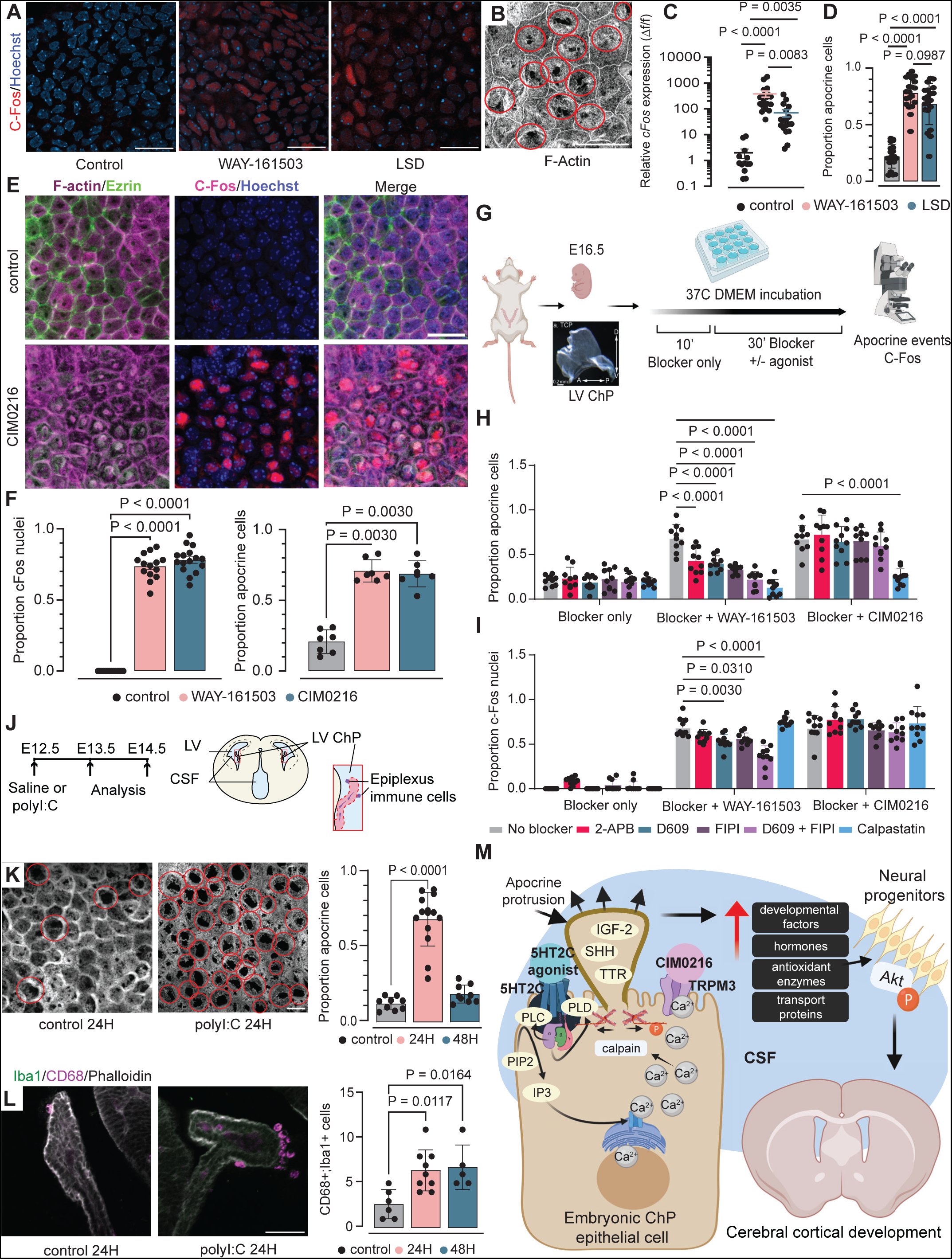
An approach for screening maternal-fetal apocrine triggers and the molecular underpinnings of apocrine secretion. (**A-B**) Confocal images of E16.5 LV ChP after maternal LSD exposure. Scale bar 20µm. (**C**) RT-PCR for *cFos* in E16.5 ChP (Saline *n* = 13, 5 litters, WAY *n* = 19, 5 litters, LSD *n =* 18, 5 litters). P values from ordinary one-way ANOVA. (**D**) Apocrine structures in E16.5 LV ChP after maternal LSD exposure. (**E**) Confocal images of E16.5 ChP after maternal saline or WAY-161503 injection. Scale bar 20µm. (**F**) Proportion of cells with c-Fos and apocrine structures in E16.5 LV ChP following maternal saline (*n* = 7, 3 litters), WAY-161503 (*n* = 7, 3 litters), or CIM0216 (*n* = 6, 3 litters) injection. P values from Mann-Whitney test. (**G**) Experiment for results in (H) and (I). (**H**) Apocrine structures or (**I**) c-Fos expression in E16.5 LV ChP after incubation with blockers and agonists. *n* = 10 explants/condition. P values from two-way ANOVA with Tukey’s multiple comparisons test. (**J**) Experiment for results in (K)-(L). (**K**) Representative confocal images and quantification of apocrine structures (circled in red) in E14.5 LV ChP after maternal saline (*n* = 9) or PolyI:C injection. (24H *n* = 14, 48H *n* = 9). P values from Mann-Whitney test. Scale bar 20µm. (**L**) Representative confocal images and quantification of epiplexus immune cells after maternal saline (*n* = 6) or PolyI:C injection (24H *n = 9*, 48H *n* = 5). Scale bar 100µm. P values from Mann-Whitney test. (**M**) Working model of ChP apocrine secretion. All data presented as means ± SEM.

Though 5HT2C activation reliably caused ChP apocrine secretion, it was not necessary for this process, as mice lacking the 5HT2C receptor still displayed the same baseline rate of apocrine secretion (**Fig. S7B-E**). Because 5HT2C is a Gαq-protein coupled receptor that induces a rapid rise in intracellular Ca^2+^, we considered whether apocrine secretion may be evoked by other processes that increase intracellular Ca^2+^ in ChP epithelial cells. We queried our ChP cell atlas (*15*) for epithelial expression of receptors associated with calcium regulation in cells. Among the most highly expressed candidate genes was *Trpm3* (*48*) (**Fig. S7F**), a heat-activated transient receptor potential cation channel subfamily M member 3 (TRPM3), which gates Na^+^ and Ca^2+^ (*49*). We pursued a straightforward approach to screen whether maternally delivered ligands of TRPM3 also alter the embryonic CSF proteome via apocrine secretion: the concurrent evaluation of apocrine structures and expression of activity-dependent protein c-Fos, an indicator of cell activation, in ChP epithelial cells. When a TRPM3 agonist, CIM0216, was administered subcutaneously to pregnant mice at E14.5, embryonic ChP apocrine structures increased more than 3-fold (**Fig. 4E-F**) and c-Fos was expressed in 73.6% of epithelial nuclei, comparable to WAY-161503 (**Fig. 4F**). This interaction may partially explain why hyperthermic episodes during pregnancy, including maternal fever or environmental factors like air temperature or hot-tub immersion, are linked to poor neurodevelopmental outcomes such as ADHD (*50*) and ASD (*51*).

Subsequent ChP explant culture experiments illuminated the necessary intracellular signaling pathway components for apocrine secretion by determining the effects of combinations of blockers with 5HT2C or TRPM3 agonists on c-Fos and apocrine structures (**Fig. 4G**). Apocrine secretion evoked via 5HT2C relied on phospholipase C, phospholipase D, and IP3 (**Fig. 4H**). Differing stimuli could lead to apocrine secretion via increased intracellular Ca^2+^, but convergence upon an intracellular pathway enabling breakdown of the actin lattice and massive cytoskeletal reorganization was necessary. We identified a convergent pathway component, calpain, that is required for both 5HT2C- and TRPM3-mediated secretion. Calpain facilitates Ca^2+^-mediated cytoskeletal remodeling by cleaving F-actin anchor protein ezrin (*52*). Inhibition of ChP calpain with calpastatin prevented formation of apocrine structures (**Fig. 4H**), though cells still responded as evidenced by c-Fos expression (**Fig. 4I**).

Apocrine secretion also occurs at supraphysiological levels in more naturalistic paradigms. Maternal immune activation (MIA) increases the risk of many neurodevelopmental disorders (*53*) and can cause cerebral cortical abnormalities (*54*), affect embryonic ChP immune cells, and create an inflammatory signature in CSF (*55*). We modelled MIA using polyinosinic-polycytidylic acid (PolyI:C) (**Fig. 4J**) and examined ChP epithelial cells for apocrine structures at 24H and 48H following maternal PolyI:C administration, revealing more than a 5-fold increase in apocrine structures at 24H (**Fig. 4K**). ChP tissue appeared to be recovering from this widespread event at 48H, with no new apocrine structures present. Propagation of MIA to the embryonic brain was apparent by increased ChP immune cells at both timepoints (**Fig. 4L**) (*55*). These findings suggest that the ChP apocrine secretion may serve to rapidly modulate CSF contents during acute inflammation. Collectively, our studies demonstrate that apocrine secretion can be evoked through various stimuli with far-reaching consequences on brain development (**Fig. 4M**).

## DISCUSSION

We report a previously uncharacterized regulatory mechanism of CSF signaling factors that are important for healthy brain development. The dominant view of ChP secretion among neuroscientists has been limited to the familiar mechanisms by which neurons release factors – exosomes and vesicles – and has not fully considered the unique capabilities of epithelial cells. As a result, it has been challenging to define physiological situations that would lead to imbalanced CSF signaling that impairs cerebral cortical development (*6*, *55*). Our data bring a more comprehensive understanding by showing that ChP epithelial cells utilize an apocrine secretion mechanism that can significantly alter the CSF proteome. It will be important for the field to consider this mechanism alongside vesicular and exosomal exocytosis when considering how the ChP regulates CSF contents.

Apocrine secretion’s functionality by the time of cerebral cortical neurogenesis underscores its critical role in early brain development. The developmental relevance of this secretion is remarkable, releasing cell-viability promoters, apoptosis inhibitors, and regulators of neuronal function and development. Mice exposed to chronic embryonic overstimulation of ChP apocrine secretion displayed a marked shift in cortical neuron populations, specifically increased Satb2+ neurons and decreased Tbr1+ neurons in the deep layers of the somatosensory cortex. This shift closely resembles what Tsyporin et al observe in mice lacking zinc-finger transcription factor *Fezf2*, which acts as a transcriptional repressor in the context of neuronal identity (*56*). This raises the intriguing possibility that CSF-borne factors serve to broadly activate or repress transcriptional pathways in neural progenitors, helping to steer them towards the proper identity. Our observations of altered deep layers further suggest that perhaps only the CSF-contacting apical progenitor cells are vulnerable to CSF alterations, since these cells are known to be the ones that migrate to form the deeper layers (*38*, *57–59*). It will be important to understand what CSF factors are exerting this transcriptional control, and through which progenitor receptors they are signaling, furthering our nuanced understanding of cerebral cortical neuron specification.

The rapidity and efficiency by which external stimuli, delivered to a pregnant mother, can activate embryonic ChP and alter CSF composition—within 30 minutes—are striking. This process, even when repeated minimally during embryonic development, profoundly impacts cortical neuron populations and adult behavior. Decreased sociability is observed across many neurodevelopmental disorders (*60*), and decreased marble burying has been observed in rodent models of long COVID (*61*). These findings have widespread implications for basic and translational neuroscience. External factors such as maternal illness or medication could influence fetal brain development through this pathway. The potential influence of substances that interact with ChP apocrine secretion pathways, like psychedelics and potentially SSRIs, on brain development is noteworthy. The ChP’s sensitivity to various stimuli and rapid accessibility also raises possibilities for targeted interventions to mitigate negative developmental impacts from these same perturbations.

ChP apocrine secretion, beyond its role in neurodevelopment, continues to occur in adult mice and can be triggered via the same stimuli and signaling pathways. This suggests potential roles for long-distance central nervous system (CNS) signaling, in which the ChP has been thought to play a role in inflammatory (*10*, *62*) and neuroendocrine (*5*) signaling. Our demonstration of a high-capacity, inducible secretion mechanism at the blood-CSF interface also raises exciting possibilities for CNS drug delivery. Given that the ChP can be targeted with AAVs, and protein production locally driven (*63*, *64*), harnessing ChP apocrine secretion may prove useful in the supplementation of CSF with health-promoting factors or pharmacological interventions.

## ACKNOWLEDGEMENTS

We thank the NIDA Drug Supply Program for providing (+)-Lysergic acid diethylamide (+)-tartrate. We thank Joel Elmquist for sharing *Htr2c-lox-STOP-lox* mice. We thank Chinfei Chen, Hisashi Umemori, Cheng-Hao Chien, Harry Cramer, and the BCH IDDRC Cellular Imaging Core for imaging assistance. We thank Maria Ericsson, Anja Nordstrom, the HMS Electron Microscopy Facility, and the NEU EM Facility for assistance with sample preparation and imaging. We thank Nathaniel Hodgson and the BCH Animal Behavior and Physiology Core for assistance with behavioral tasks and analysis. We thank Towia Libermann, Simon Dillon, and the BIDMC Genomics, Proteomics, Bioinformatics and Systems Biology Center for assistance with SomaScan. We thank members of the Lehtinen laboratory, Mark Andermann, Jeffrey Macklis, Pascal Kaeser, Timothy Hla, and Nancy Chamberlin for discussion and/or comments on the manuscript. We acknowledge BioRender.com for assistance with creating several schematics.

## AUTHOR CONTRIBUTIONS

Conceptualization: YC, ND, MKL. Methodology: YC, JPH, ND, FBS, MKL. Performed experiments: YC, JPH, EDY, ND, FBS. Data curation and analysis: YC, TAL. Supervision and funding: MKL. Writing – original draft: YC, MKL. Writing – review & editing: All authors.

## DATA AND MATERIALS AVAILABILITY

All data are available in the manuscript or through the supplementary materials.

## MATERIALS AND METHODS

### Mice

The Boston Children’s Hospital IACUC approved all experiments involving mice in this study. Adult and timed-pregnant CD1 male and female mice were obtained from Charles River Laboratories. Animals were housed in a temperature- and humidity-controlled room (70 ± 3 °F, 35–70% humidity) on a 12 h light/12 h dark cycle (7 a.m. on 7 p.m. off) and had free access to food and water.

### Injection of WAY-161503, LSD

All agonists were injected subcutaneously into timed-pregnant CD1 female mice in a vehicle of 0.9% sterile saline solution. Dosages were as follows: WAY-161503 3mg/kg (Tocris 1801), (+)-Lysergic acid diethylamide tartrate 0.2 mg/kg (NIDA Drug Supply Program).

### Behavioral analyses

All behavioral analyses were carried out in collaboration with the Boston Children’s Hospital Animal Behavior & Physiology Core.

#### Three chamber social approach

Social approach was tested in a modified automated three-chambered apparatus using methods similar to those previously described (*65–67*). Noldus EthoVision XT videotracking software (version 9.0, Noldus Information Technologies, Leesburg, VA) was employed to increase throughput. The updated apparatus was a rectangular three-chambered box, 40 × 60 × 23 cm, fabricated from matte white acrylic (P95 White, Tap Plastics, Sacramento, CA). Opaque retractable doors (12 × 33 cm) were designed to create optimal entryways between chambers (5 × 10 cm), while providing maximal division of compartments. Three zones, defined using the EthoVision XT software, detected time in each chamber for each phase of the assay. Zones to score sniffing were defined as the annulus extending 2 cm from each novel object or novel mouse enclosure (inverted wire cup, Galaxy stainless steel pencil and utility cup, Kitchen Plus, http://www.kitchen-plus.com). Direction of the head, facing toward the cup enclosure, defined sniff time. A top mounted infrared-sensitive camera (Ikegami ICD-49, B&H Photo, New York, NY) was positioned directly above every two three-chambered units. Infrared lighting (Nightvisionexperts.com) provided uniform, low level illumination.

The subject mouse was first placed in the center chamber for 10 min, then allowed to explore all three empty chambers during a 10-min habituation session, then allowed to explore the three chambers with a novel object in one side chamber and a novel mouse in the other side chamber. Lack of innate side preference was confirmed during the initial 10 min of habituation to the entire arena. Novel stimulus mice were 129Sv/ImJ, a relatively inactive strain, aged 10– 14 weeks old, and were matched to the subject mice by sex. Number of entries into the side chambers served as a within-task control for levels of general exploratory locomotion. Sociability in this assay is defined as more time in the chamber with the novel mouse than in the chamber with the novel object, and more time sniffing the novel mouse than sniffing the novel object, within each genotype or within each treatment group.

#### Male-female ultrasonic vocalizations

All mice were tested in a 33 × 15 × 14 cm plastic cage with 3 cm of sawdust and a metal flat cover. Male experimental subjects were habituated to this apparatus for 30 min prior to testing; an unfamiliar stimulus female mouse (an adult NMRI female, different for each tested male) was then introduced into the testing cage and left there for 3 min. Previous studies have shown that in these experimental settings USVs are emitted only by the male mouse in the male-female interaction (*68*, *69*). Furthermore, all spectrograms obtained here were additionally inspected to exclude the presence of “double calls,” i.e., overlapping in their timing, but with different, non-harmonic, characteristics (e.g., different peak and mean frequency) which would indicate the concomitant emission of USVs by the two interacting subjects during testing.

Testing sessions were recorded by a camera placed on the side of the cage and videos analyzed with Observer XT (Noldus, Netherlands). One observer who was unaware of the genotype and stress conditions of the animals scored the behavior of the test male mice, quantifying the time spent performing affiliative behaviors (*70–72*) i.e., sniffing the head and the snout of the partner, its anogenital region, or any other part of the body; contact with partner through traversing the partner’s body by crawling over/under from one side to the other or allogrooming. Non-social activities were also measured: rearing (standing on the hind limbs sometimes with the forelimbs against the walls of the cage); digging; self-grooming (the animal licks and mouths its own fur).

An ultrasonic microphone UltraSoundGate Condenser Microphone CM 16 (Avisoft Bioacoustics, Berlin, Germany) was mounted 2 cm above the cover of the testing cage; it was connected via an UltraSoundGate 116 USB audio device (Avisoft Bioacoustics) to a personal computer, where acoustic data were recorded with a sampling rate of 250 kHz in 16-bit format by Avisoft Recorder (version 2.97; Avisoft Bioacoustics).

Recordings were then transferred to the Sonotrack Call Classification Software (version 1.4.7, Metris B.V., Netherlands). This fully automatic software recognizes different USV types and calculates quantitative parameters including the total number and mean duration of the calls.

Based on previous literature on call types (*43*, *73–75*), the following USV types were selected for automatic recognition in our dataset: Short, Flat, (Ramp) Up, (Ramp) Down, Chevron, Step-Up, Step-Down, Step-Double (Split), Complex-3, Complex-4, Complex-5, Complex-5+.

#### Marble burying

Clean cages (27×16.5×12.5 cm) were filled with 4.5-cm corncob bedding, gently overlayed with 20 black glass marbles (15 mm diameter) equidistant in a 4×5 arrangement. Testing consisted of a 30-min exploration period. The number of marbles buried (>66% marble covered by bedding material) was recorded. White noise (55 dB) was present during testing. Light intensity was held at 30 lux.

### ChP tissue dissection

Whole embryonic and adult ChP tissues were isolated as follows. LV ChP: Cerebral cortical hemispheres were separated. Following incisions into the cortical tissue at either end of the hippocampus, the hippocampus was rolled out using the flat surface of a microscalpel and the underlying LV ChP was separated from the hippocampus/fornix. 3V ChP: The dorsal midline of the midbrain was exposed and 3V ChP was isolated. Specific to adult brain, flushing the midline region with 1xHBSS helped visualize the ChP, which was identifiable by a blood vessel running along its rostro-caudal axis. The adult 3V ChP extends ventrally and rostrally, ultimately connecting to the ventral base of the LV ChP. 4V ChP: Hindbrain was separated from the brain. The cisterna magna was exposed by gently guiding away the developing or mature cerebellum to expose the 4V ChP.

### Transmission Electron Microscopy

LV ChP was micro-dissected in HBSS (Thermo Fisher Scientific) and drop-fixed immediately in FGP fixative (5% Glutaraldehyde, 2.5% Paraformaldehyde and 0.06% picric acid in 0.2 M sodium cacodylate buffer, pH 7.4). After 2 hours at RT, the tissue was washed in 0.1M cacodylate buffer and postfixed with 1% Osmium tetroxide (OsO4)/1.5% Potassium ferrocyanide (KFeCN6) for 1 hour, washed twice in water, washed once in 50 mM Maleate buffer pH 5.15 (MB), incubated in 1% uranyl acetate in MB for 1 hour followed by one wash in MB, 2 washes in water, and subsequently dehydrated in an series of increasingly concentrated alcohol (10 min each; 50%, 70%, 90%, 2×10 min 100%). The samples were immersed in propyleneoxide for 1 hour and infiltrated ON in a 1:1 mixture of propyleneoxide and TAAB Epon (TAAB Laboratories Equipment Ltd, https://taab.co.uk). The following day, the samples were embedded in TAAB Epon and polymerized at 60°C for 48 hours. Ultrathin sections (about 80 nm) were cut on a Reichert Ultracut-S microtome and placed on copper grids stained with lead citrate. Images were acquired with a JEOL 1200EX transmission electron microscope and recorded with an AMT 2k CCD camera (Electron Microscopy Facility, Harvard Medical School).

### Scanning Electron Microscopy

Lateral ventricle ChP tissue from adult and embryonic mice was micro-dissected and post-fixed in 1.0% osmium tetroxide in 0.1M cacodylate buffer (pH 7.4) for 1 hour at room temperature. Following postfixation, the samples were rinsed with buffer then dehydrated through a graded series of ethanol. The specimens were then critical point dried with CO2 using a Samdri PVT-3 critical point dryer (Tousimis Corp. Rockville, MD). The specimens were attached to specimen mounts using conductive adhesive tabs, coated with 5nm platinum using a Cressington 208HR sputter coater (Cressington Scientific Instruments, Ltd. Walford, UK). Imaging was conducted on a Hitachi S-4700 FESEM. Electron Microscopy Imaging, consultation and services were performed in the HMS Electron Microscopy Facility and the NEU Electron Microscopy Facility.

### Hematoxylin and Eosin (H&E) Staining

Paraffin-embedded brains were sectioned to a thickness of 5 mm. The sections were deparaffinized in xylene and then rehydrated via successive incubations in 100% ethanol, 95% ethanol, and water. Sections were incubated in Gill 3 hematoxylin (Sigma Aldrich, St. Louis, MO) for 2 min., followed by a 5s incubation in 0.5% ammonia water to increase the contrast of the hematoxylin stain. Next, sections were rinsed in water and incubated in 1% alcoholic eosin for 3 min. Finally, sections were dehydrated via successive incubations in 95% ethanol and 100% ethanol and mounted onto glass slides with Permount (Thermo Fisher Scientific).

### Immunostaining

*Choroid plexus explants.* ChP explants were dissected and fixed in 4% paraformaldehyde for 10 min at room temperature. Samples were incubated in primary antibodies overnight at 4°C, shaking and in secondary antibodies at room temperature for two hours, shaking. All antibodies were diluted in 0.1% Triton X-100 in PBS. No blocking or antigen retrieval methods were used. All samples were counterstained with Hoechst 33342 (Invitrogen, H3570, 1:10,000) and mounted onto slides using Fluoromount-G (SouthernBiotech).

*Brain sections.* Postnatal animals were perfused with PBS followed by 4% paraformaldehyde (PFA). The brains were quickly dissected and post-fixed with 4% PFA overnight. Samples were cryoprotected and prepared for embedding at 4°C with 10% sucrose, 20% sucrose, 30% sucrose, 1:1 mixture of 30% sucrose and OCT (overnight), and OCT (1 h on ice). Samples were frozen in OCT. Antigen retrieval and blocking were not performed. Cryosections were permeabilized (0.1% Triton X-100 in PBS), incubated in primary antibodies at 4°C overnight and secondary antibodies for 2 h at room temperature. Hoechst 33,342 (Invitrogen H3570, 1:10,000, 5 min at room temperature) was used to visualize nuclei. Sections were mounted on glass slides with Fluoromount-G (SouthernBiotech). Images were acquired using Zeiss LSM980 confocal microscope with 20× objective. ZEN Black software was used for image acquisition and ZEN Blue used for Airy processing.

### Immunohistochemical and immunoblotting antibodies

#### Primary Antibodies

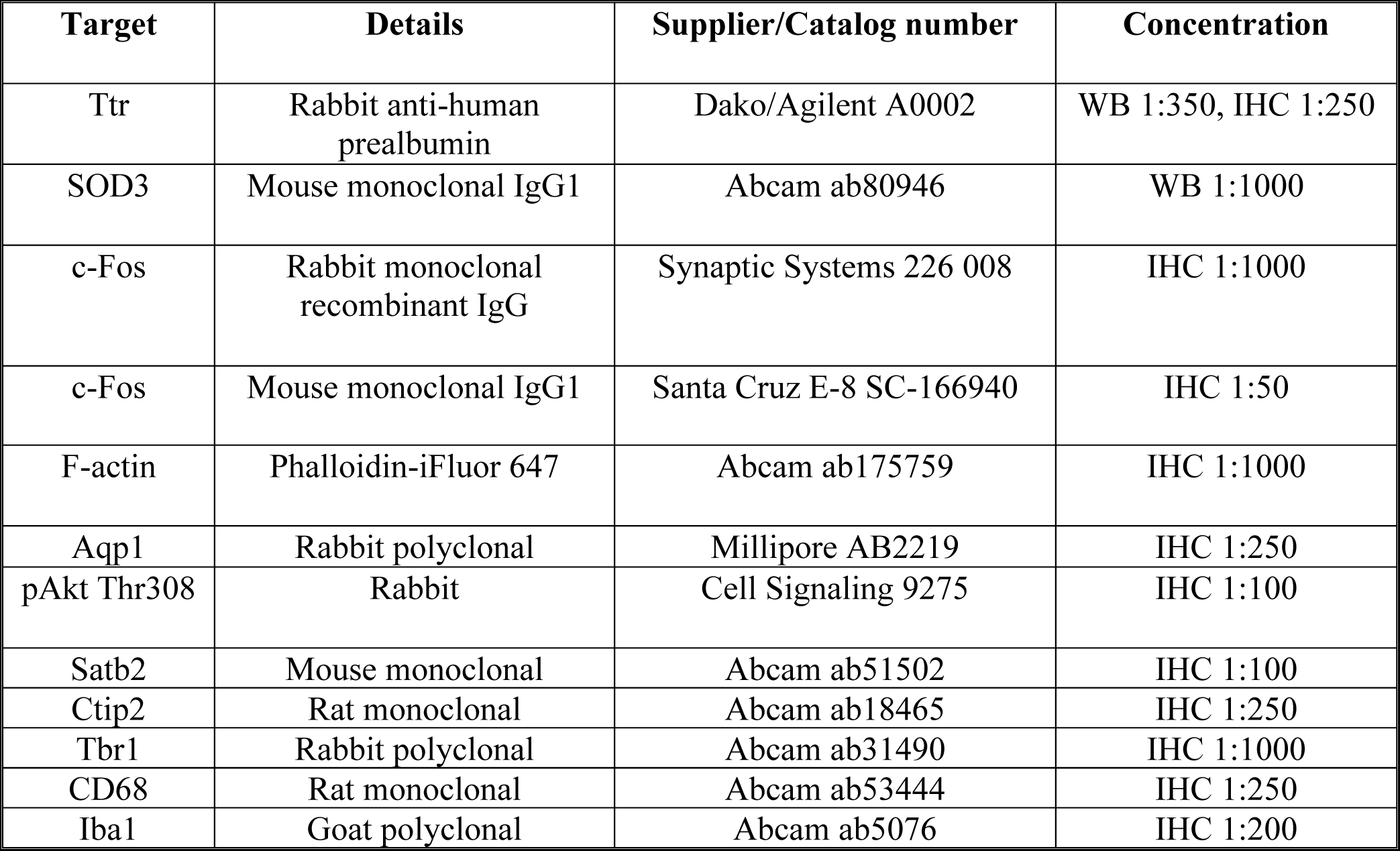

| Target | Details | Supplier/Catalog number | Concentration |
| --- | --- | --- | --- |
| Ttr | Rabbit anti-human prealbumin | Dako/Agilent A0002 | WB 1:350, IHC 1:250 |
| SOD3 | Mouse monoclonal IgG1 | Abcam ab80946 | WB 1:1000 |
| c-Fos | Rabbit monoclonal recombinant IgG | Synaptic Systems 226 008 | IHC 1:1000 |
| c-Fos | Mouse monoclonal IgG1 | Santa Cruz E-8 SC-166940 | IHC 1:50 |
| F-actin | Phalloidin-iFluor 647 | Abcam ab175759 | IHC 1:1000 |
| Aqp1 | Rabbit polyclonal | Millipore AB2219 | IHC 1:250 |
| pAkt Thr308 | Rabbit | Cell Signaling 9275 | IHC 1:100 |
| Satb2 | Mouse monoclonal | Abcam ab51502 | IHC 1:100 |
| Ctip2 | Rat monoclonal | Abcam ab18465 | IHC 1:250 |
| Tbr1 | Rabbit polyclonal | Abcam ab31490 | IHC 1:1000 |
| CD68 | Rat monoclonal | Abcam ab53444 | IHC 1:250 |
| Iba1 | Goat polyclonal | Abcam ab5076 | IHC 1:200 |

#### Secondary Antibodies

| Host | Target | Name | Supplier/Catalog number | Concentration |
| --- | --- | --- | --- | --- |
| Donkey | Goat | Alexa fluor 488 | Abcam AB96931 | IHC 1:250 |
| Goat | Mouse | Alexa fluor 594 | Life Technologies A-11005 | IHC 1:250 |
| Goat | Mouse | Alexa fluor 647 | Thermo Fisher A-21235 | IHC 1:250 |
| Goat | Rabbit | Alexa fluor 488 | Life Technologies A11034 | IHC 1:250 |
| Goat | Rabbit | Alexa fluor 594 | Abcam a11012 | IHC 1:250 |
| Goat | Rat | Alexa fluor 568 | Life Technologies A-11077 | IHC 1:250 |
| Goat | Rabbit | HRP | Life Technologies 31460 | WB 1:1000 |
| Goat | Mouse | HRP | Life Technologies 31430 | WB 1:1000 |

### Image acquisition and quantification

Immunohistochemical images were acquired using either a Zeiss LSM710 or LSM980 microscope at the Boston Children’s Hospital Cellular Imaging Core. Images of ChP explants and brain sections were acquired using a 20x air or 63x oil objective. Images of ChP explants prepared with protein-retention expansion microscopy were acquired with a 63x NA1.2 water-immersion objective. ZEN Black software was used for image acquisition and ZEN Blue used for Airy processing.

### Fluorescence intensity quantification

Fluorescence intensity quantification (Fig. 1E, Fig. 2M) was performed in ImageJ/FIJI. The desired region was outlined with the freehand ROI tool. Desired parameters were set by going to Analyze > Set Measurements and selecting Area, Integrated Density, and Mean Grey Value. Analyze > Measure was used to acquire initial intensity value. Then, 3 small areas of the image with no fluorescence were selected to be background. Analyze > Measure was used to acquire background intensity value. Corrected total region fluorescence was calculated as Integrated Density – (Area of Selected Region x Mean Fluorescence of Background readings).

### Apocrine structure quantification

Apocrine structure quantification was performed by hand by author YC. YC took 4 20x fields of view per ChP explant and counted epithelial cells displaying apocrine structures via either SEM or IHC. The proportion of cells with apocrine structures was quantified as the number of epithelial cells with an apocrine protrusion divided by the total number of visible epithelial cells. The proportion of cells with apocrine structures for each of the 4 fields of view was then averaged and used as N=1 biological replicate per embryo.

### C-Fos positive nuclei quantification

C-Fos positive nuclei quantification was performed in ImageJ/FIJI. 4 20x fields of view were taken per ChP explant, and the average proportion of C-Fos positive nuclei in these 4 fields of view was used as N=1 biological replicate per embryo. Each field of view was first thresholded, then the “Analyze particles” function used with a size discrimination of over 200 px. This was performed for C-Fos channels and DAPI channels, and then the number of C-Fos nuclei divided by the total number of nuclei to obtain “Proportion of C-Fos positive nuclei”.

### Excitatory neuronal marker counting and co-expression analysis

Cerebral cortical layer marker counting and colocalization was performed using Biodock software (*76*). A fully automated AI model was trained by a 3-step iterative process on 3 test staining images containing Ctip2, Sat2, Tbr1, and DAPI. Input images were 4 channels. TIF files obtained by importing the .CZI file from the microscope into FIJI, rotating, and cropping a rectangular region comprising the primary somatosensory cortex from just above layer 1 to the corpus callosum.

The AI model was trained to classify each cell in the cerebral cortex into one of 8 classes based on co-expression of each marker used. Potential cell classes were:

1. ctip2- satb2- tbr1- (DAPI only)
2. ctip2+, satb2-, tbr1-
3. ctip2+, satb2+, tbr1-
4. ctip2+, satb2+, tbr1+
5. ctip2+, satb2-, tbr1+
6. ctip2-, satb2+, tbr1-
7. ctip2-, satb2+, tbr1+
8. ctip2-, satb2-, tbr1+

Each image’s AI analysis was carefully checked by author YC, validating the correct identification of cells and visualizing overall distribution of labels.

The data resulting from the Biodock AI model is a .CSV file containing a comprehensive list of cell measurements obtained from cortical tissue samples. The data included several parameters for each cell: Object ID, Image Origin, Class, Area, Eccentricity, Length of Major and Minor Axes, Perimeter, Solidity, X and Y Positions, and Average Intensity across four channels.

Further analysis was conducted in the R programming environment. We utilized key packages including ggplot2 (*77*) for data visualization, dplyr (*78*) for data manipulation, and tidyr (*79*) for data tidying. A critical step in our analysis involved segmenting the cortical area into 6 equal-sized bins from top to bottom. To achieve this, we calculated the minimum and maximum values of the Y Position from our data, establishing six equally spaced bins along the Y-axis. This segmentation allowed us to analyze cell distribution across different cortical layers. Each cell was assigned to one of the six bins based on its Y Position, facilitating a layer-specific analysis of cell distribution. We then performed a detailed count of cells in each bin, categorizing them into various classes based on their expression of markers: ctip2, satb2, and tbr1.

We computed the percentage of each cell class in each bin, relative to the total cell count in that bin. The final step was exporting our analyzed data into a new CSV file, named based on the original file, for ease of reference and further analysis. This file contained the expanded summary table with cell counts, percentages, and other relevant statistics. These data were imported into GraphPad Prism for statistical analyses and graphing.

In the interest of scientific transparency and reproducibility, the complete R script used for this analysis is available at https://doi.org/10.5281/zenodo.10464093.

### Real-Time Quantitative PCR (RT-qPCR)

Choroid plexus tissues were rapidly microdissected from the lateral and fourth ventricles of mouse brains submerged in cold PBS. Tissues were immediately frozen on dry ice and stored at −80C. RNA samples were prepared using the Monarch Total RNA Miniprep Kit (New England Biolabs #T2010S) for total RNA isolation, following the manufacturer’s instructions and eluting in 50uL nuclease-free water. Each biological replicate consisted of two paired lateral ventricle ChP explants or one 4^th^ ventricle explant. The extracted RNA was quantified using a spectrophotometer, and 100 ng of RNA was reverse-transcribed into cDNA using the LunaScript RT Master Mix Kit (New England Biolabs #E3025L). qPCR reactions utilized specific primers and were performed in triplicate with Applied Biosystems TaqMan Gene Expression Master Mix (ThermoFisher #4369016). Probes used were Mm00487425_m1 (Fos, FAM-MGB, ThermoFisher #4331182) and Hs99999901_s1 (Eukaryotic 18s rRNA, VIC-MGB, ThermoFisher #4331182). The cycling process was carried out using the Roche LightCycler 480II. Relative gene expression analysis was conducted using the 2^(-ΔΔCT) method (*80*), normalizing expression levels to 18S rRNA as the internal control.

### Immunoblotting

Embryonic CSF was collected and centrifuged at 10000G for 5 minutes, then flash-frozen on dry ice and stored at −80°C. Samples were denatured in 2% SDS supplemented with 2- mercaptoethanol by heating at 95°C for 5 min. Equal amounts of CSF were loaded and separated by electrophoresis in a NuPAGE 4–12% Bis-Tris gel (Invitrogen #NP0322), transferred to a nitrocellulose membrane (250 mA, 1.5 h, on ice), blocked in filtered 5% milk in TBST, incubated with primary antibodies overnight at 4°C followed by HRP conjugated secondary antibodies (1:1,000) for 2h, and visualized with ECL or ECL select substrate.

*Silver stain.* Silver staining was performed on a NuPAGE 4–12% Bis-Tris gel (Invitrogen #NP0322) loaded with 0.5µL of embryonic CSF per lane. After electrophoresis, silver staining was performed using the Invitrogen SilverQuest Staining Kit (Invitrogen #LC6070) according to manufacturer instructions.

### Protein-retention expansion microscopy

Protein-retention expansion microscopy was carried out using established protocols (*81*). Importantly, we found that of many we tried, only sodium acrylate purchased from Combi-Blocks (QC-1489, 98% purity, CAS number 7446-81-3, FW 94.0, Batch B41013) was sufficiently pure for the preparation to work. Images were acquired in Boston Children’s Hospital Cellular Imaging Core using a Zeiss LSM710 confocal microscope with a 63x 1.2 NA water immersion objective.

### Expansion Microscopy Reagents

| Name | Supplier/Catalog number |
| --- | --- |
| Acryloyl-X SE 5mg | Thermo Fisher Scientific A20770 |
| Proteinase K buffered aqueous glycerol solution, for molecular biology, >800 units/mL | Sigma P4850-5ML |
| Ammonium Persulfate 100g | Sigma A3678-100G |
| 4-Hydroxy-TEMPO 5g | Sigma 176141-5G |
| Acrylamide 500g | Sigma A9099-500G |
| N,N'-methylenebisacrylamide 100g | Sigma 146072-100G |
| Poly-L-Lysine solution 0.1% in H<sub>2</sub>O, 100mL | Millipore sigma P8920-100mL |
| Sodium acrylate 25g | Combi-Blocks QC-1489 |
| Anhydrous DMSO | Sigma 276855-100ML |

### Proteomics

Cerebrospinal fluid was collected from embryonic mice at gestational day E16.5 A fine glass capillary was inserted into the cisterna magna to withdraw CSF, and samples were only included if they did not contain any blood or tissue contamination. CSF from 5 litters of embryos was pooled for each sample, as the SomaScan assay used requires 40uL of CSF, and each E16.5 embryo yields ~2-3 uL of CSF. Data reflects 3 biological replicates (pooled samples) each for a vehicle-injected control condition and a WAY-161503 injection condition. Collected CSF samples were immediately flash frozen on dry ice and transferred to −80°C for storage to preserve protein integrity.

Samples were then transferred to the Beth Israel Deaconess Medical Center Genomics, Proteomics, Bioinformatics and Systems Biology Center, which performed proteomic profiling using the oligonucleotide aptamer-based proteomics technology, SomaScan (*82*), provided by SomaLogic (Boulder, CO; SomaScan Assay v4.1) according to the manufacturer’s standardized protocol. SomaScan Assay v4.1 uses 7322 high affinity, distinct aptamer reagents to measure expression of 6596 unique proteins. SomaLogic’s proprietary protein-capture reagents called SOMAmers (Slow Off-rate Modified Aptamer) leverages Cy3 fluorescently-labeled, chemically modified, short single-stranded DNA sequences known as aptamers that form three-dimensional structures similar to antibodies and based on their unique nucleotide sequences are able to bind with high affinity and selectivity to distinct proteins. The readout for the assay are custom micro-array chips, containing complementary oligonucleotides to each individual fluorescently labeled SOMAmer. Three provided kit controls and one no-protein buffer control were analyzed in parallel with the CSF samples to adjust for intra-run variation. Median normalization and calibration of the SomaScan data were performed according to standard quality control (QC) protocols at SomaLogic. The final assay readout is in relative fluorescence units (RFU), an arbitrary unit directly proportional to the protein expression.

### Proteomics analysis - R

#### Data Acquisition and Preprocessing

The normalized dataset for this study was obtained from SomaLogic and stored in the .adat format. The file, named “original.adat,” was processed using a script written in R (version 4.1.2). The readat package was utilized to import the raw data (*83*). Sample classification was performed based on the experimental design, with samples grouped as Saline, WAY, or Quality Control. Quality Control samples were subsequently removed from the dataset.

#### Data Analysis and Visualization

A suite of R packages including reshape2 (*84*), magrittr (*85*), dplyr (*78*), ggplot2 (*77*), limma (*86*), Biobase (*87*), countdata (*88*), ggrepel (*89*), topGO (*90*), org.Hs.eg.db (*91*), glue (*92*), tibble (*93*), Rgraphviz (*94*), enrichplot (*95*), DOSE (*96*) and ggvenn (*97*) were employed for data analysis and visualization. The limma package was used for linear regression and analysis, while countdata facilitated fold change analysis.

Proteins of interest were identified based on the largest between-group variation, and their intensity levels were visualized using ggplot2. Differential expression analysis was performed, and results were visualized as volcano plots. Gene names were standardized using the org.Hs.eg.db package, which provides genome-wide annotation for human genes.

#### Gene Ontology Enrichment Analysis

The topGO package was used to perform Gene Ontology (GO) enrichment analysis (*90*). This analysis focused on identifying significantly enriched GO terms within the list of differentially expressed genes, using Fisher’s exact test and the Kolmogorov-Smirnov test. The GO terms were categorized into three main ontologies: Biological Process (BP), Molecular Function (MF), and Cellular Component (CC).

#### Statistical Analysis

Statistical significance for differential expression was determined using a combination of fold change and t-test p-values. For GO enrichment analysis, significance was assessed using the Fisher’s exact test and the Kolmogorov-Smirnov test, as implemented in the package. Adjusted p-values were calculated to control for multiple testing.

#### Data Visualization

Data visualization was extensively performed using ggplot2, Rgraphviz, enrichplot, and ggvenn. This included the generation of volcano plots for differential expression analysis, heatmaps for visualizing gene expression patterns, and Venn diagrams to illustrate the overlap between different protein sets. Gene Ontology terms were visualized using ggplot2 and ggpubr (*98*), focusing on terms with a significant number of genes and highlighting the statistical significance using a star-based system.

In the interest of scientific transparency and reproducibility, the complete R script used for this analysis is available at https://doi.org/10.5281/zenodo.10464124.

### Proteomics analysis – Ingenuity pathway analysis

For pathway analysis, the Ingenuity Pathway Analysis (IPA) software (QIAGEN Inc.) was employed (*36*). IPA is a web-based software application used for the integration, interpretation, and analysis of data derived from genomics, transcriptomics, proteomics, and metabolomics studies. It helps in identifying new targets or candidate biomarkers within the context of biological systems. Once differentially abundant proteins were identified in R, a list of up- and down- regulated proteins and their expression values was used for Ingenuity Pathway Analysis.

#### Importing data

The normalized and preprocessed SomaScan data were imported into IPA. The data file included Entrez Gene Symbol identifiers for each protein along with their corresponding fold change values. Data were formatted to meet IPA requirements, ensuring correct mapping of protein identifiers to IPA’s knowledge base.

#### Core Analysis

IPA’s core analysis was performed, which included the identification of significantly enriched pathways, upstream regulators, molecular networks, and biological functions. The analysis settings were adjusted to the specific needs of the study, including the selection of appropriate species (mouse), tissue types, and cell types, as relevant.

#### Pathway and Network Identification

IPA identified relevant pathways that were significantly enriched in the dataset, providing insights into the biological functions most prominently represented. Molecular interaction networks were generated, highlighting key proteins and the interactions among them within the context of the identified pathways.

#### Upstream Regulator Analysis

Upstream regulator analysis in IPA was used to predict potential upstream molecules that could be responsible for the observed changes in protein expression. This analysis also helped in understanding the cause-and-effect relationship between upstream regulators and downstream effects in the dataset. IPA provided statistical scores for the enrichment of pathways and networks based on the input data. The significance of the association between the dataset and canonical pathways was measured in terms of p-values, calculated using Fisher’s exact test.

#### Data Interpretation and Visualization

IPA offered various tools for the interpretation and visualization of results. This included graphical representations of pathways, networks, and their components, with options to customize views for detailed analysis. The results were interpreted in the context of current biological knowledge and relevant literature integrated within IPA’s database. The analysis was performed on a standard computing setup with internet access to utilize the web-based IPA software. No additional computational resources were required as the processing and analysis were handled by IPA’s servers.

### Explant conditioned media experiment

Choroid plexus explants were rapidly microdissected from embryonic brains and separated by ventricle. Upon dissection, they were placed in 50 microliters of oxygenated artificial CSF as previously described (*19*) in a 96-well plate, on ice. 5 embryonic explants were pooled per well. For figure 2D-E, explants were incubated with 5µM of WAY-161503 (5HT2C agonist) or with a vehicle control for 30 minutes at 37°C. Then, the conditioned aCSF was removed, flash-frozen on dry ice, and stored at −80°C. For figure 4E-F, explants were again grouped 5 per well in 50µL of oxygenated artificial CSF. Explants were incubated for 10 minutes at 37°C with any applicable blockers, and then introduced to WAY-161503 (5HT2C agonist) or CIM0216 (TRPM3 agonist). Explants were incubated with these agonists for another 30 minutes at 37°C, then explants were processed for immunohistochemistry as described above.

### Blocker and agonist information was as follows

**Blocker and agonist information was as follows:**
| Compound | Supplier | Target | Concentration |
| --- | --- | --- | --- |
| 2-APB | RND Systems 1224 | IP3 receptor antagonist | 50 mM |
| D609 | RND Systems 1437 | PC-PLC inhibitor | 50 mM |
| FIPI | RND Systems 3600 | PLD inhibitor | 50 mM |
| Calpastatin | Millipore sigma 208902 | Calpain I and II inhibitor | 10 µM |
| WAY-161503 | Tocris 1801 | 5HT2C agonist | 1 mM |
| CIM0216 | Tocris 5521 | TRPM3 agonist | 5 µM |
| Serotonin hydrochloride | Sigma-Aldrich H9523 | Serotonin receptors | 5 µM |

### ELISAs

ELISA assays were performed with embryonic CSF that was flash-frozen on dry ice and stored at −80°C after collection. IGF-II was measured with the R&D Systems Mouse/Rat/Porcine/Canine IGF-II/IGF2 Quantikine ELISA Kit (MG200) according to manufacturer guidelines. SHH was measured with the R&D Systems Mouse Sonic Hedgehog/Shh N-Terminus Quantikine ELISA Kit MSHH00 according to manufacturer guidelines.

### Maternal immune activation

For timed pregnancies, females were checked for the presence of plugs, and the date of the plug was noted as embryonic day 0.5 (E0.5). Body weight was monitored every 3–4 days, and ultrasound was available to help confirm gestational age. On E12.5, pregnant dams received a single dose (20 mg/kg, i.p.) of polyinosinic-polycytidylic acid (polyI:C, Sigma Aldrich) or saline as vehicle control.

### Quantification and statistical analysis

To achieve robust and unbiased results, we performed our analyses across multiple samples, we prospectively randomized treatment group via random number generation, quantified results depending on genotype in a double-blinded manner and performed rigorous pre- and post-hoc statistical analyses to test our hypotheses. We consulted regularly with statisticians at Harvard (BCH, BIDMC, Broad) to ensure proper calculations of power analyses and statistical methods including accounting for multiple comparisons and covariates. We performed an equal number of experiments on males and females and carefully examined sex as a biological variable in our analyses. For each experiment, pilot studies were used to determine variance, and two-tailed power analyses determined optimal group size for alpha = 0.05 and beta = 0.80. Biological replicates (*N*) were defined as samples from distinct individual animals, analyzed either in the same experiment or within multiple experiments, with the exception when individual animal could not provide sufficient sample (i.e., CSF), in which case multiple animals were pooled into one biological replicate and the details are stated in the corresponding figure legends.

Statistical analyses were performed using Prism (GraphPad) v7 or R (version 3.6.3). Outliers were excluded using ROUT method (*Q* = 1%). Appropriate statistical tests were selected based on the distribution of data, homogeneity of variances, and sample size. *F* tests or Bartlett’s tests were used to assess homogeneity of variances between datasets. Parametric tests (*T* test, ANOVA) were used only if data were normally distributed, and variances were approximately equal. Otherwise, nonparametric alternatives were chosen. Analyses were done as specified in figure captions using One-way ANOVA, Student’s two-tailed unpaired *t* test (when *F* test failed to discover unequal variances), or Welch’s two-tailed unpaired *t* test (when *F* test discovered unequal variances) if data followed a parametric distribution. If data were non-parametric, Kruskal-Wallis test or Mann-Whitney test was used. Data are presented as means ± standard deviation (SD). If multiple measurements were taken from a single individual, data are presented as means ± standard errors of the mean (SEMs). Please refer to figure legends for sample size. *p* values < 0.05 were considered significant. Exact *P* values are marked in the figures where space allows.

