## Supplemental Material for "A choroid plexus apocrine secretion mechanism shapes CSF proteome and embryonic brain development"

**The PDF file includes:**

Figures S1 to S7
Tables S1 to S3

**Other Supplementary Materials for this manuscript include the following:**

Movies S1 to S3
Data S1

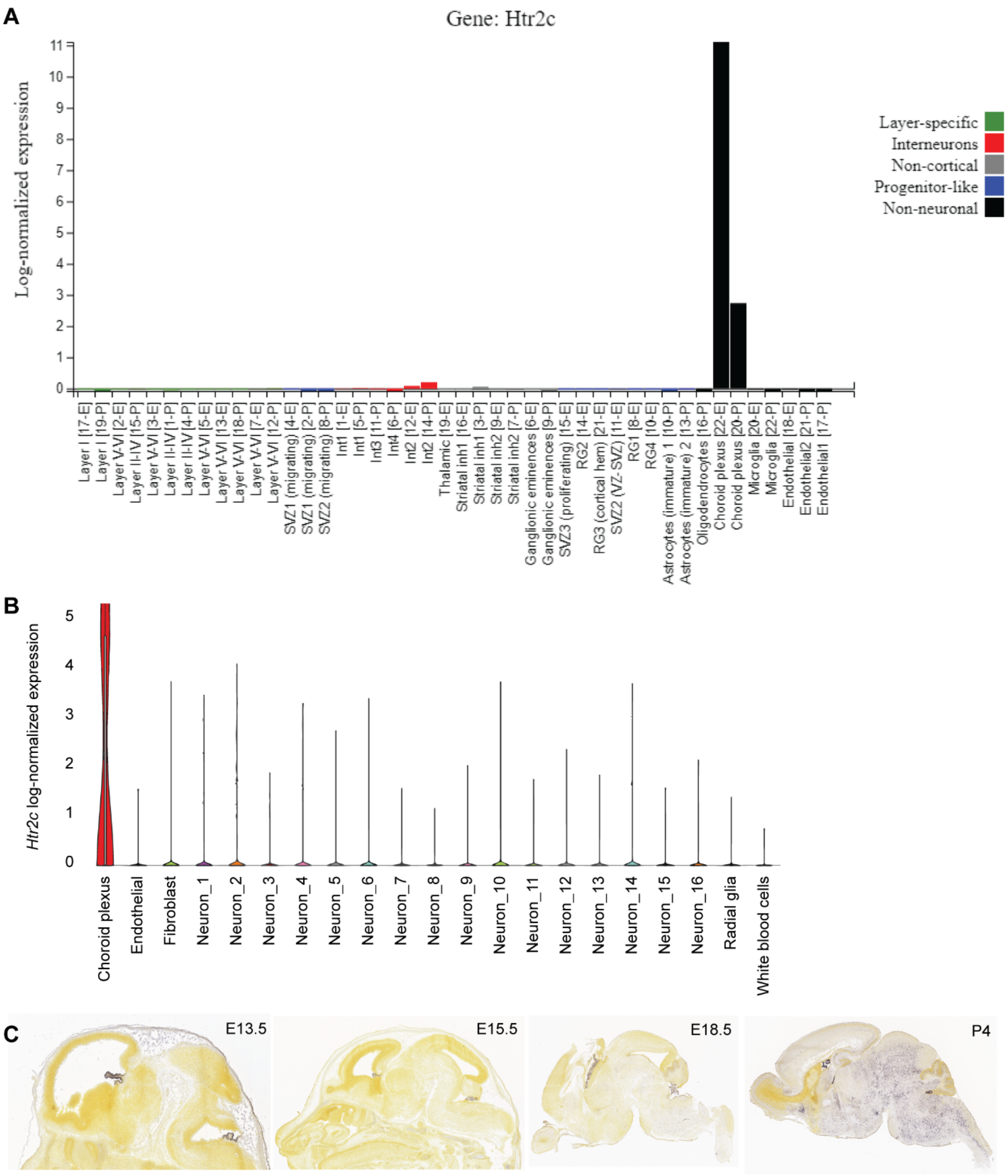
A
Gene: Htr2c
Log-normalized expression
0
1
2
3
4
5
6
7
8
9
10
11
Layer-specific
Interneurons
Non-cortical
Progenitor-like
Non-neuronal
Layer I [17-E]
Layer I [19-P]
Layer V-VI [2-E]
Layer II-IV [15-P]
Layer V-VI [3-E]
Layer II-IV [1-P]
Layer II-IV [4-P]
Layer V-VI [5-E]
Layer V-VI [13-E]
Layer V-VI [18-P]
Layer V-VI [7-E]
Layer V-VI [12-P]
SVZ1 (migrating) [4-E]
SVZ1 (migrating) [2-P]
SVZ2 (migrating) [8-P]
Int1 [1-E]
Int1 [5-P]
Int3 [11-P]
Int4 [6-P]
Int2 [12-E]
Int2 [14-P]
Thalamic [19-E]
Striatal inh1 [16-E]
Striatal inh1 [3-P]
Striatal inh2 [9-E]
Striatal inh2 [7-P]
Ganglionic eminences [6-E]
Ganglionic eminences [9-P]
SVZ3 (proliferating) [15-E]
RG2 [14-E]
RG3 (cortical hem) [21-E]
SVZ2 (VZ-SVZ) [11-E]
RG1 [8-E]
RG4 [10-E]
Astrocytes (immature) 1 [10-P]
Astrocytes (immature) 2 [13-P]
Oligodendrocytes [16-P]
Choroid plexus [22-E]
Choroid plexus [20-P]
Microglia [20-E]
Microglia [22-P]
Endothelial [18-E]
Endothelial2 [21-P]
Endothelial1 [17-P]
B
Htr2c log-normalized expression
0
1
2
3
4
5
Choroid plexus
Endothelial
Fibroblast
Neuron_1
Neuron_2
Neuron_3
Neuron_4
Neuron_5
Neuron_6
Neuron_7
Neuron_8
Neuron_9
Neuron_10
Neuron_11
Neuron_12
Neuron_13
Neuron_14
Neuron_15
Neuron_16
Radial glia
White blood cells
C
E13.5
E15.5
E18.5
P4

**Figure S1. In the embryonic brain, *Htr2c* is selectively expressed in ChP**. (A) Bar graph of E14.5 and P0 *Htr2c* expression reveals that, throughout the mouse neocortex, ChP selectively expresses during embryonic development. Notably, *Htr2c* is not expressed in progenitors. Created with the Zylka Lab's Single-cell transcriptomic analysis of mouse neocortical development gene-by-gene search webpage with data from Loo et al 2019 (*38*). https://zylkalab.org/datamousecortex (B) Violin plot of E14 *Htr2c* expression demonstrates ChP selectivity of *Htr2c* expression in mouse brain at E14.5. created with Broad Institute Single Cell Portal from Mouse E14 brain single nucleus RNA sequencing data gathered for Russell et al 2023 (*38*). https://singlecell.broadinstitute.org/single_cell/study/SCP2170/slide-tags-snrna-seq-on-mouse-embryonic-e14-brain (C) Expression of *Htr2c* in E13.5, E15.5, E18.5, and P4 mouse brain shows that before E18.5, *Htr2c* is exclusively expressed in ChP. At E18.5, low expression arises in brainstem nuclei. At P4, *Htr2c* expression is widespread. Allen Developing Mouse Brain Atlas, developingmouse.brain-map.org/experiment/show/100047619.

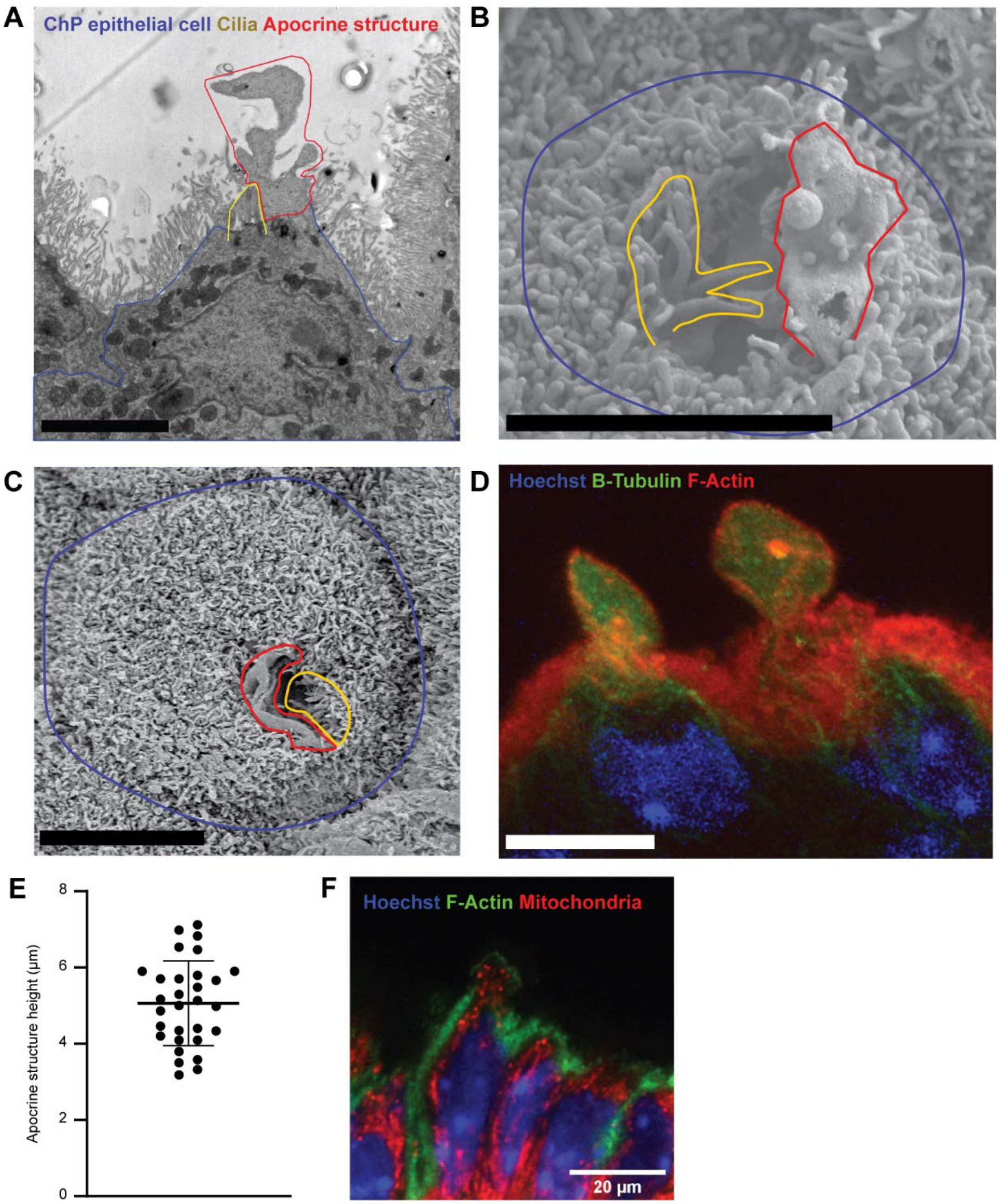
A
ChP epithelial cell Cilia Apocrine structure
B
C
D
Hoechst B-Tubulin F-Actin
E
Apocrine structure height (µm)
0
2
4
6
8
F
Hoechst F-Actin Mitochondria
20 µm

**Figure S2. Choroid plexus apocrine structures.** All scale bars 5μm. (A) Transmission electron microscopy image depicting an E16.5 ChP epithelial cell with an apocrine structure protruding immediately adjacent to cilia. (B) Scanning electron microscopy (SEM) image depicting an E16.5 ChP epithelial cell with a collapsed, likely post-secretion apocrine structure protruding immediately adjacent to cilia. (C) SEM image depicting an E16.5 epithelial cell with an emerging apocrine structure protruding immediately adjacent to cilia. (D) Expansion microscopy image depicting two E16.5 ChP epithelial cells, each with apocrine structures supported by a network of microtubules. (E) Quantification of size of apocrine structures at E16.5, N = 30 mice from 6 litters. (F) Expansion microscopy image depicting ChP LV epithelial cells, one with an apocrine protrusion containing mitochondria (Citrate Synthase antibody).

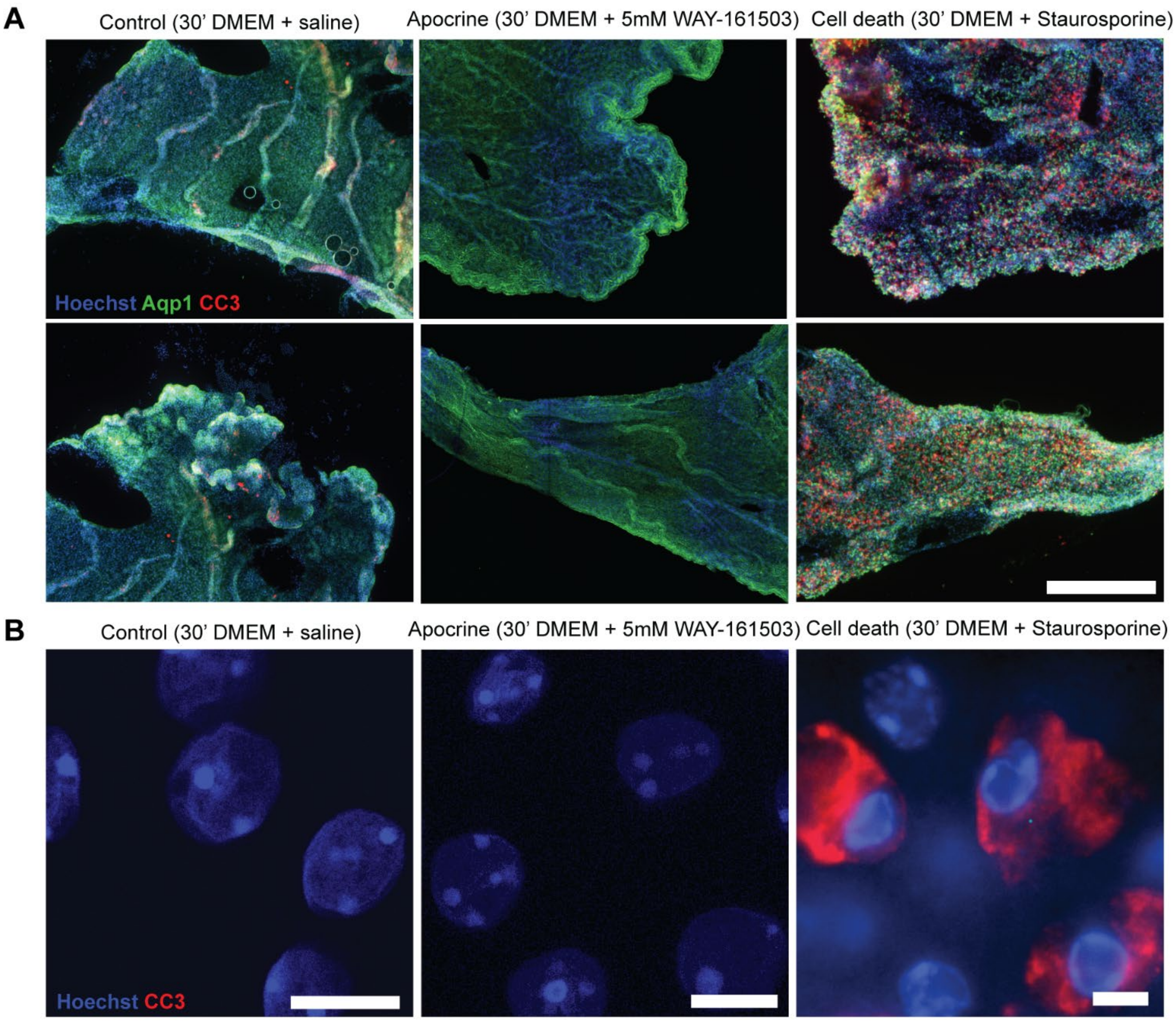


**Figure S3. Choroid plexus apocrine secretion is not apoptotic.** (A) Representative confocal images of LV ChP explants, scale bar 100µM. CC3 staining reveals apoptosis in a staurosporine-treated positive control for cell death (right), but not after evoking apocrine secretion (middle). (B) Representative confocal images of LV ChP nuclei reveal apoptosis and nuclear chromatin condensation in positive control for cell death (right) but not after apocrine secretion (middle). Scale bars all 5µM.

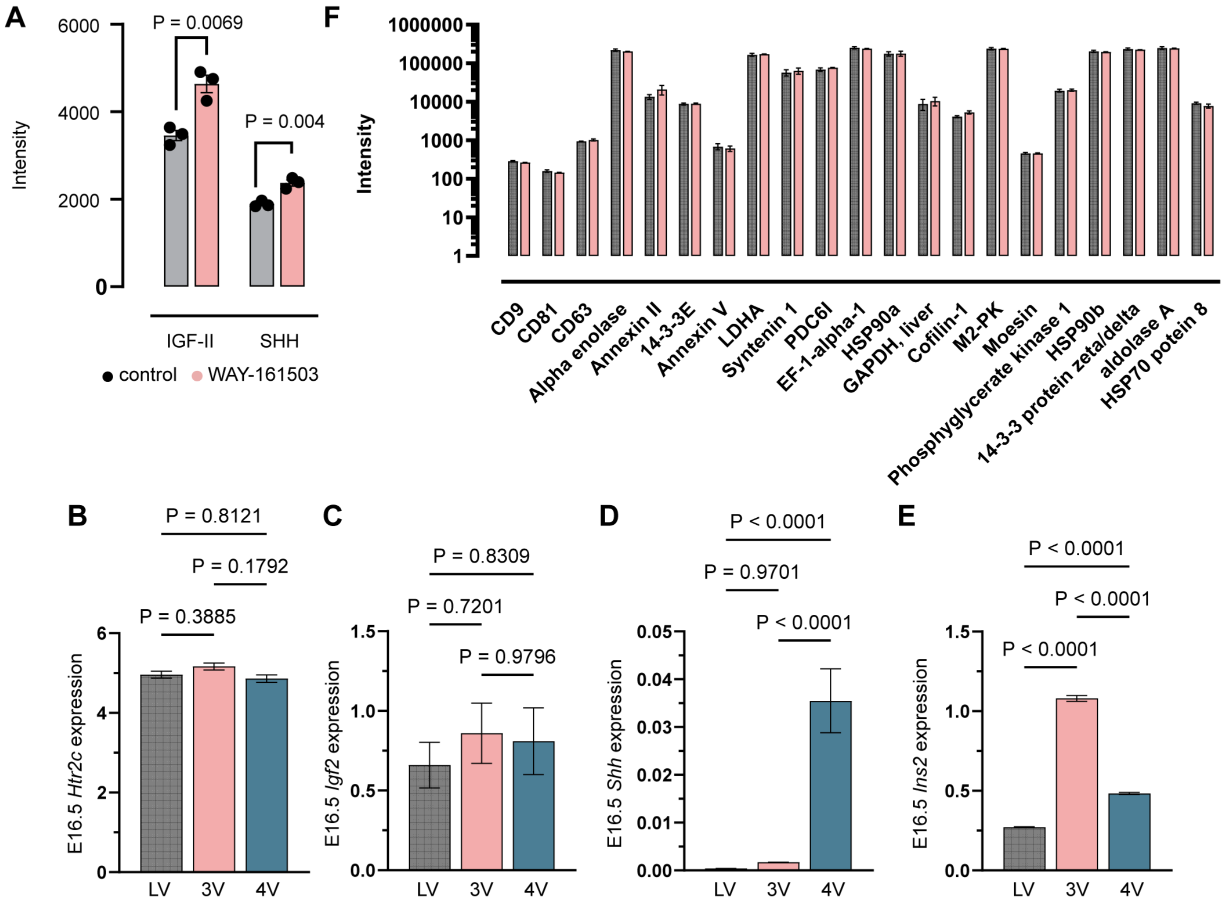


**Figure S4. ChP apocrine proteomics and regionalized gene expression**. (A) SomaScan intensity measurements for IGF-II and SHH protein in CSF demonstrate apocrine release. N = 3 E16.5 CSF samples, each pooled from 3 litters. (B) *Htr2c* (C) *Igf2* (D) *Shh* and (E) *Ins2* expression across ventricles in E16.5 ChP epithelial cells. All values from Dani *et al* 2020. (F) SomaScan intensity measurements for a list of exosomal proteins show no significant increases in CSF after apocrine secretion.

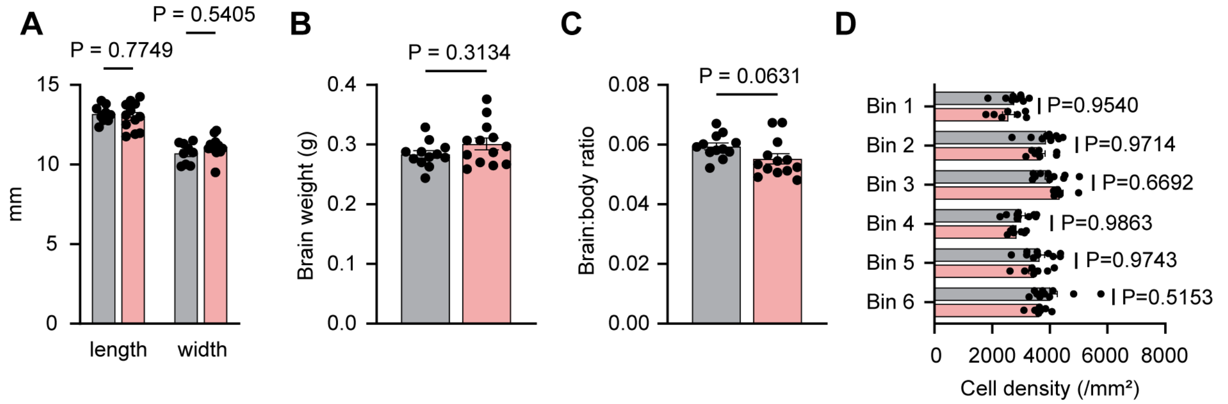


**Figure S5. Chronic embryonic overstimulation of apocrine secretion does not lead to gross morphological brain changes.** (A) P30 brain dimensions in control (*n* = 12) and chronic apocrine (*n* = 13) mice. P values from Mann-Whitney test. (B) Quantification of P30 brain weight and (C) Brain:body weight ratio in control (*n* = 12) and apocrine (*n* = 13) mice. P values from Mann-Whitney test. (D) Cell density in P8 S1 cortex across 6 bins in control (*n* = 6) and apocrine (*n* = 10) mice. P values from Mann-Whitney test.

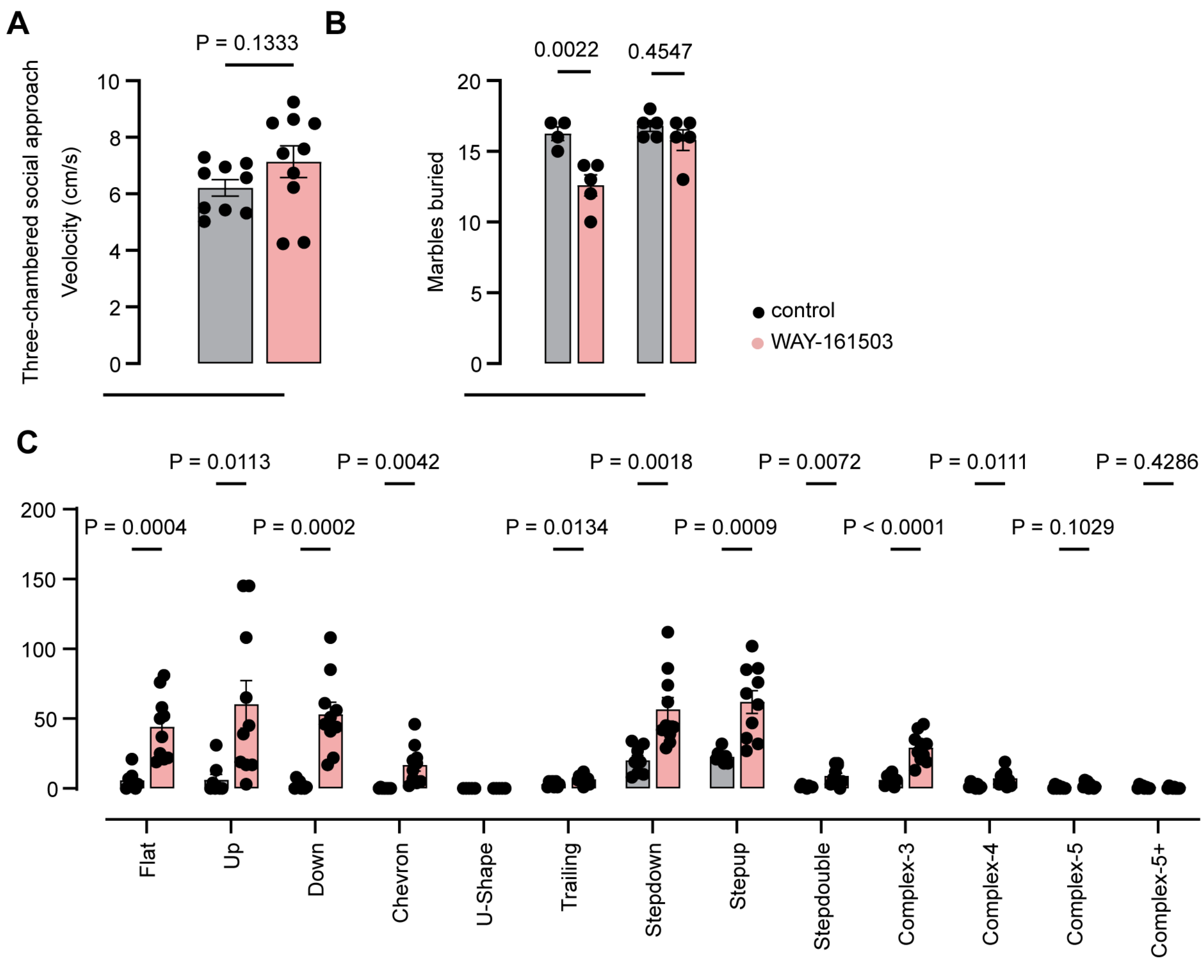


**Figure S6. Embryonic overstimulation of apocrine secretion leads to adult behavioral changes.** (A) Velocity in three-chambered social approach task is not significantly influenced by embryonic apocrine secretion. (B) Adult male mice bury fewer marbles after embryonic overstimulation of apocrine secretion, adult female mice are unaffected. (C) Many subtypes of adult male-female ultrasonic vocalizations increase after embryonic overstimulation of apocrine secretion.

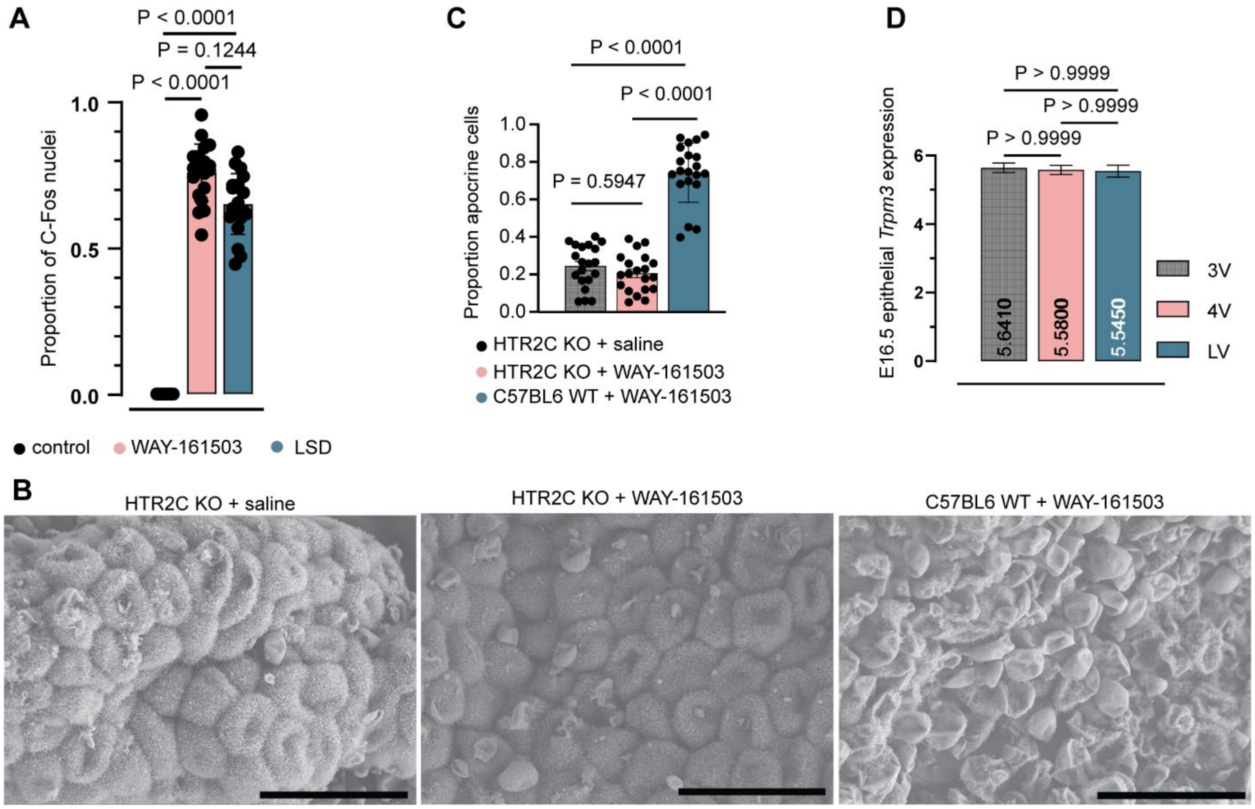


**Figure S7. HTR2C receptor is not necessary for baseline rates of apocrine secretion**. All scale bars 30µm. (A) Representative SEM of E16.5 HTR2C KO mouse LV ChP 30' after maternal injection of a saline vehicle control. (B) Representative SEM of E16.5 HTR2C KO mouse LV ChP 30' after maternal injection of WAY-161503. (C) Representative SEM of WT C57BL6 E16.5 mouse LV ChP 30' after maternal injection of WAY-161503. (D) Quantification of apocrine structures. N = 20 mice from 4 litters. Comparisons performed with ordinary one-way ANOVA. (E) Quantification of c-Fos positive nuclei in E16.5 ChP 30' after maternal saline, WAY-161503, or LSD administration. (F) *Trpm3* expression across ventricles in E16.5 ChP epithelial cells.

| Protein | Fold Change | Protein | Fold Change | Protein | Fold Change |
|---|---|---|---|---|---|
| CFL2 | -2.677702396 | ANGPT1 | 1.107722 | CAPN2 | 1.253244 |
| SLMAP | -1.958047138 | APBB1 | 1.122756 | SHH | 1.254541 |
| MGMT | -1.489150925 | EIF3B | 1.135808 | CHMP3 | 1.256712 |
| NPM1 | -1.465645541 | EDAR | 1.140927 | SULT1B1 | 1.259806 |
| ACBD4 | -1.3126736 | TYRP1 | 1.144411 | HSD17B10 | 1.263076 |
| FAM171A1 | -1.306032518 | YWHAH.1 | 1.14687 | GFRA1 | 1.263377 |
| HSPA5 | -1.289718269 | NTN1 | 1.149025 | COL2A1.1 | 1.264269 |
| FIBP | -1.239313573 | NTN1.1 | 1.149307 | TLL1 | 1.27252 |
| CPTP.1 | -1.216096043 | MIF | 1.15044 | PPP2R1A | 1.273816 |
| RBM39 | -1.199984307 | CADM1 | 1.156866 | EFNA2 | 1.2745 |
| CYTL1 | -1.196602888 | MAP2K3 | 1.16222 | DMKN | 1.287389 |
| LILRB5 | -1.187896776 | SYAP1 | 1.162807 | CHST1 | 1.287997 |
| MYBPC2 | -1.187287086 | ENTPD6 | 1.163364 | EIF4B | 1.30313 |
| TNFRSF25 | -1.152109912 | DNAJC18 | 1.164101 | NEGR1 | 1.315286 |
| YTHDC1 | -1.13757003 | TPST2 | 1.165798 | FGFRL1.1 | 1.329058 |
| NOSIP | -1.134353999 | FAM171B.1 | 1.166307 | IGF2 | 1.340103 |
| ACVR1 | -1.13010118 | STK3 | 1.17156 | GLT8D1 | 1.340113 |
| MET.1 | -1.111471542 | POMC.2 | 1.173689 | EFNA5 | 1.348281 |
| TMEM9 | -1.094764252 | NFASC | 1.175271 | NLGN2.1 | 1.351096 |
| IGHG2 | -1.093790708 | UBXN4.1 | 1.184758 | THSD7A | 1.352122 |
| CLUH | -1.085820113 | ACLY | 1.189366 | NEGR1.1 | 1.354892 |
| LRP11.1 | -1.079009834 | ANAPC10 | 1.189717 | NUCB1 | 1.363485 |
| THAP2 | 1.073949919 | CMPK1 | 1.193424 | C11orf68 | 1.368757 |
| WBP2 | 1.074126956 | PTPRU | 1.196621 | OTUD7B | 1.372287 |
| SHBG.1 | 1.07450517 | EPN1 | 1.200221 | VAPA | 1.390194 |
| TMEM52 | 1.076830491 | SLAMF1 | 1.208397 | SBDS | 1.395874 |
| DGKB | 1.077596117 | HAUS1 | 1.211578 | NUCB2 | 1.400142 |
| MAGEA3 | 1.081612965 | SIGLEC11.1 | 1.212659 | NLGN3.1 | 1.439247 |
| CGA\|LHB | 1.082951889 | ING1 | 1.216631 | ACAT2 | 1.463587 |
| EPHB3 | 1.083444976 | SERPINE2 | 1.224738 | LDHA.1 | 1.467336 |
| ULBP3 | 1.08347554 | MCTS1 | 1.225412 | KREMEN1 | 1.468277 |
| SKP1 | 1.085563306 | TPST1 | 1.22785 | IVD | 1.497916 |
| DFFA | 1.089705016 | HOMER2 | 1.229924 | PTK7 | 1.617312 |
| FLRT1.1 | 1.089984573 | PARK7.2 | 1.23303 | DSTN | 1.751027 |
| DDI1 | 1.092664998 | PCDH9 | 1.238185 | SLITRK6 | 1.975016 |
| UBE2D3.1 | 1.094902212 | NDRG3 | 1.24016 | TYMS | 1.998942 |
| NRG1.1 | 1.095929315 | ANXA6 | 1.242997 | AMIGO2 | 2.336661 |
| NTN4 | 1.100043446 | GALNT14 | 1.246098 | AMIGO2.1 | 2.453852 |
| CHST15 | 1.102332036 | HSPH1 | 1.246523 | NLGN4Y | 2.932438 |
| ATP23 | 1.10241375 | FTCD | 1.253056 | NLGN3 | 3.033497 |
| PDGFA | 1.104117096 | | | | |
| CD2AP | 1.106712066 | | | | |
| GRIA4 | 1.10749003 | | | | |

**Table S1.** Protein list showing differential abundance by SomaLogic analysis. All differentially abundant proteins between control CSF and apocrine CSF at E16.5.

| Protein | SomaScan Target Name |
|---|---|
| HSPA8 | HSP70 protein 8 |
| CD9 | CD9 |
| GAPDH | GAPDH, liver |
| ACTB | not assayed |
| CD63 | CD63 |
| CD81 | CD81 |
| ANXA2 | annexin II |
| ENO1 | Alpha enolase |
| HSP90A1 | HSP 90a |
| EEF1A1 | EF-1-alpha-1 |
| PKM | M2-PK |
| YWHAE | 14-3-3E |
| SDCBP | Syntenin 1 |
| PDC6IP | PDC6I |
| ALB | not assayed |
| YHWAZ | 14-3-3 protein zeta/delta |
| EEF2 | not assayed |
| ACTG1 | not assayed |
| LDHA | LDHA |
| HSP90AB1 | HSP 90b |
| ALDOA | aldolase A |
| MSN | Moesin |
| ANXA5 | Annexin V |
| PGK1 | phosphoglycerate kinase 1 |
| CFL1 | Cofilin-1 |

**Table S2.** List of top 25 proteins that are often identified in exosomes from http://exocarta.org/exosome_markers_new and their SomaScan target name identifier. "Not assayed" means an aptamer for this protein was not included in the 7000-plex SomaScan assay.

| | | |
|---|---|---|
| A1BG | GLRX | PIP |
| ACTN4 | GPX3 | PKM |
| ADCYAP1 | GSTP1 | PLA2G2A |
| ALDH1A1 | HP | PSAP |
| ANXA2 | HSP90B1 | PSMA7 |
| APOB | HSPA5 | PSMB1 |
| APOH | HSPG2 | RNASET2 |
| ARG1 | IGHM | RPS19 |
| AZGP1 | IL36G | RRBP1 |
| B2M | KLK7 | SBSN |
| CA2 | LDHA | SCGB1D1 |
| CALR | LGALS3 | SCGB2A1 |
| CAT | LPO | SCPEP1 |
| CP | LTF | SERPINA12 |
| CPA4 | LYZ | SFRP1 |
| CST1 | MDH1 | SH3GLB1 |
| CST2 | MDH2 | SORT1 |
| CST4 | MSLN | SPINT1 |
| CSTB | NEU1 | STAT3 |
| CTSB | NUCB1 | TALDO1 |
| CTSD | NUCB2 | TOLLIP |
| DMBT1 | PDCD6IP | TPM3 |
| ECHDC1 | PDIA4 | VCL |
| ENO1 | PEBP1 | WFDC2 |
| ERP29 | PFN1 | ZG16B |
| FTH1 | PIGR | |

**Table S3. Apocrine proteins identified in E16.5 CSF.** 77 of the 100 "canonical biomarkers of apocrine secretion" identified in **(14)** are present in baseline mouse CSF via SomaScan at embryonic day E16.5.

**Movie S1.** Expansion microscopy of ChP epithelial cell with an apocrine structure shows juxtaposition to cilia. Movie depicts moving through Z planes in space, “upwards”, from a cross section of the nucleus of the epithelial cell to the cell surface and then through a protruding apocrine structure. Hoechst in blue, B-Tubulin in red, Aqp-1 in green.

**Movie S2.** Expansion microscopy of ChP epithelial cells shows a network of microtubules underneath apocrine structures. Movie depicts a rotating 3D rendering of two cells with apocrine structures. Hoechst in blue, B-tubulin in green, Aqp-1 in magenta.

**Movie S3.** Expansion microscopy of ChP epithelial cells reveals TTR inside apocrine structures. Movie depicts a rotating 3D rendering of several ChP epithelial cells with apocrine structures. Hoechst in blue, F-actin in magenta, Aqp-1 in green, TTR in red.

**Data S1.** (separate file) SomaScan proteomics raw data. .adat file containing raw SomaScan data for baseline vs. apocrine E16.5 CSF. Also contains buffer controls, mixed baseline and apocrine CSF controls, and calibration samples. https://doi.org/10.5281/zenodo.10465931
